# Microglia Adopt Temporally Specific States after Irradiation, Correlating with Neuronal Asynchrony

**DOI:** 10.1101/2025.02.15.638339

**Authors:** Alejandro Lastra Romero, Efthalia Preka, Giusy Pizzirusso, Luis Enrique Arroyo-García, Georgios Alkis Zisiadis, Nuria Oliva-Vilarnau, Yana Ruchiy, Thea Seitz, Kai Zhou, Arturo G. Isla, Lara Friess, Ying Sun, Maria Querol Canut, Alia Shamikh, Yiran Xu, Changlian Zhu, Carlos F. D. Rodrigues, André Fisahn, Bertrand Joseph, Lena-Maria Carlson, Adamantia Fragkopoulou, Volker M Lauschke, Christer Betsholtz, Ahmed M Osman, Klas Blomgren

## Abstract

Cranial radiotherapy causes progressive neurocognitive impairments in cancer survivors. Neuroinflammation is a key contributor, but its dynamics and consequences for brain function remain poorly understood. Here, we performed comprehensive longitudinal profiling from 6 hours to 1 year after irradiation (IR) of the mouse hippocampus, using transcriptomic, protein, and histological analyses. We identified delayed microglial responses initiated by mitotic progression coupled interferon signaling. IR rewired the parenchymal phagocyte profiles, triggered by progressive microglial loss, failure of repopulation through self-renewal, and compensatory generation of microglia-like cells derived from peripheral monocytes. These findings were also observed in autopsied human brain. Finally, we demonstrate two phases of neuronal asynchrony, an early one associated with inflammation and a late one associated with aberrant synaptic regulation. These results provide comprehensive, longitudinal insights into microglia responses that can aid in tailoring therapies to preserve cognition in cancer survivors.

## Introduction

Cranial irradiation (IR) is standard of care in the treatment of high-grade primary and metastatic brain tumors. However, it causes long-term neurocognitive complications in 50-90% of the patients, especially in children ^1,2^. Impaired cognitive domains include learning, processing speed, memory, executive function, and attention ^3^. Attempts to omit upfront craniospinal IR in brain cancers with good prognosis aiming to reduce IR-associated neurotoxicity, by replacing it with focal IR, have failed ^4^. IR-induced cognitive deficits are, thus, an unavoidable, debilitating clinical problem that demands careful attention, particularly since the overall survival rates and remaining life expectancies of children with brain tumors by far exceed those of adults, and are expected to improve even further with current advances in cancer therapies ^5^.

The molecular underpinnings of IR-induced cognitive deficits remain largely unknown. For the past two decades, a dominant hypothesis was the depletion of postnatal hippocampal neurogenesis, a process by which new neurons are continuously generated from dividing neural stem and progenitor cells (NSPCs) in the subgranular zone (SGZ), and important for maintaining learning and memory throughout life ^6–9^. IR-induced depletion of neurogenesis has been linked to an increase in the number and activation status of microglia within the neurogenic zone, creating a hostile microenvironment that hinders NSPC proliferation and neuronal differentiation ^10,11^. Despite the compelling evidence linking reduced neurogenesis to neuroinflammation ^12^, the negative effects of inflammation on brain functions and its consequent cognitive deficits are unlikely limited to depletion of hippocampal neurogenesis, as evident from other CNS disease models ^13^.

Microglia, the resident immune cells and phagocytes in the brain, play key roles in neuroinflammation ^14^. Microglia respond rapidly to IR and undergo a series of molecular events, resulting in the production of cytokines and chemokines and the engulfment of dying NSPCs ^15–17^. There is a fundamental lack of defined mechanisms by which the IR-induced microglial responses contribute to cognitive deficits. Current knowledge is based on *in vitro* or *in vivo* studies using only one or a few time points close to the time of IR, using conventional histological analyses and measurements of pre-selected inflammatory mediators in the brain, reviewed in reference ^18^. Available data based on bulk or single-cell transcriptomic analyses of post-IR microglial responses are limited to time points within days to weeks post-IR ^16,19–21^, resulting in a need for high-resolution information about the delayed molecular events in microglia. Given the dynamic nature of microglia ^22,23^, and that post-IR cognitive deficits are delayed and progress over 5-10 years ^1^, it is imperative to investigate trajectories at the cellular and tissue level over a relevant period of time.

We performed unbiased, longitudinal *in vivo* studies of the post-IR inflammatory response in the hippocampus, a brain structure central to cognition ^24^, spanning acute and chronic phases (6 hours to 1 year), using transcriptomics combined with histological and protein analyses. We uncovered delayed microglial ‘reactivation’ and induction of unique temporal states yielding multifaceted inflammatory profiles.

## Results

### Irradiation causes biphasic inflammatory waves in the hippocampus

We have previously shown that hippocampal microglia in the juvenile brain are activated as early as 2 h post-IR and return to baseline within 1 week (wk) ^16^. However, data from adult rodent models suggest that IR induces persistent microglial activation and chronic neuroinflammation ^12,25,26^. Thus, we asked whether long-term inflammation occurs also in the juvenile brain and, if so, what role(s) microglia play. To this end, we first performed a longitudinal unbiased bulk RNA sequencing (RNA-seq) analysis of hippocampal tissue from 6 h to 6 wk post-IR. Juvenile mice were subjected to whole-brain irradiation (WBI) with a single dose of 8 Gy. This IR dose is equivalent to a total radiation dose of 18 Gy when delivered in repeated 2-Gy fractions, as in a clinical setting, estimated using the linear-quadratic (LQ) model and an α/β ratio of three for late effects in normal brain tissue ^27^.

Hippocampi were dissected 6 h (acute phase), 1 day (acute phase), 1 wk (early sub-acute phase), 2 wk (delayed sub-acute phase), and 6 wk (sub-chronic phase) post-IR, and from age-matched sham controls (SH) (Figure 1A). At all time points, principal component analysis (PCA) showed that IR samples were distinctly clustered separate from their respective SH samples (Figure 1B), indicating long-term transcriptomic alterations in the hippocampus post-IR. Numerous differentially expressed genes (DEGs; *q* < 0.05) between the SH and IR animals were detected as early as 6 h post-IR (Figure 1C). Downregulated DEGs peaked 1 day post-IR and then decreased over time (Figure 1C; Supplementary list 1). The dynamics of upregulated DEGs were different. The number of upregulated DEGs peaked 6 h post-IR, decreased by 1 day and 1 wk, followed by a second increase at 2 wk, and a decrease again 6 wk post-IR (Figure 1C; Supplementary list 1), suggesting a second delayed response 2 wk post-IR. While the earlier alterations have been extensively studied ^11,15–17,28^, we focused our attention on this later response 2 wk post-IR.

**Figure 1:**
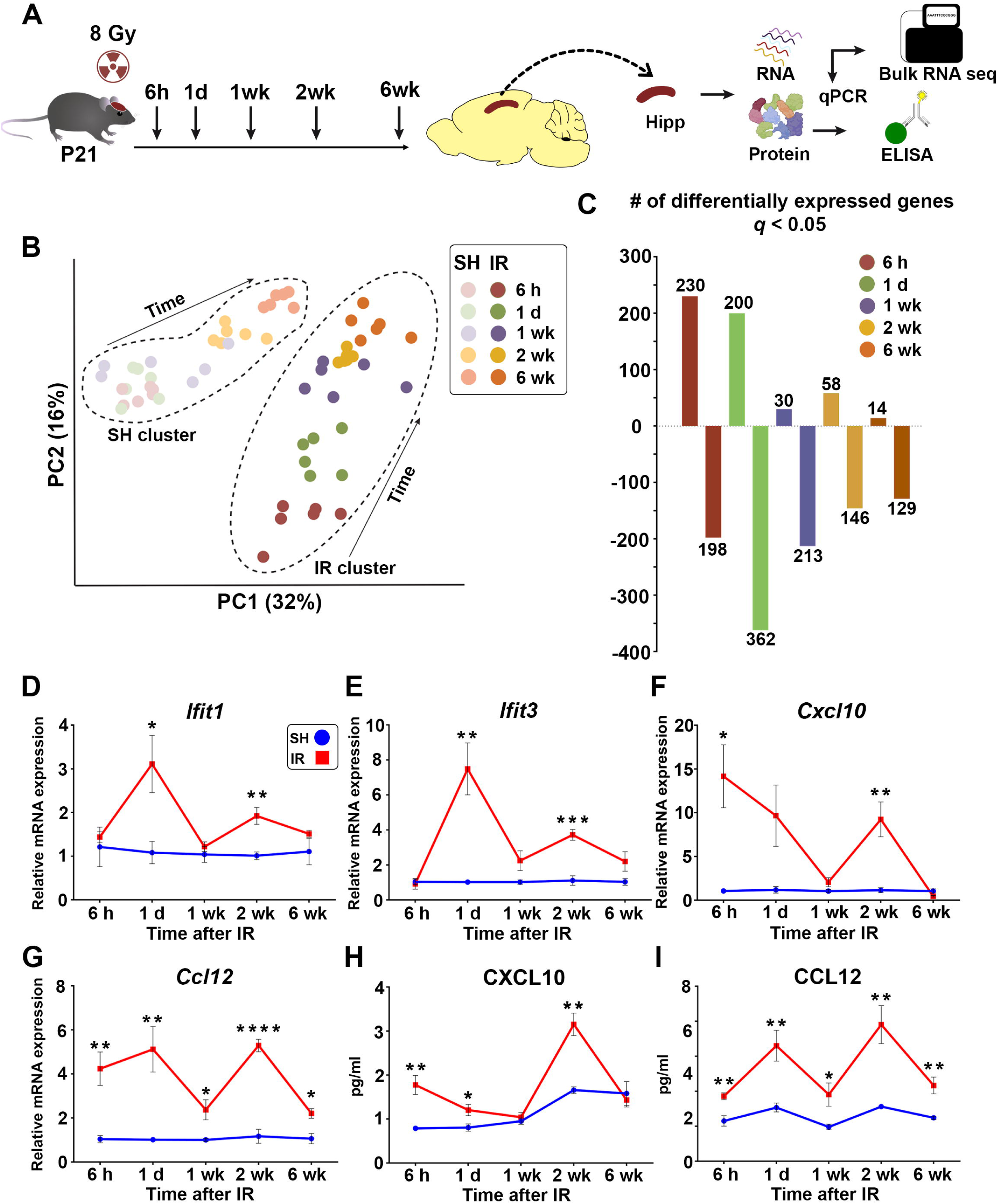
Irradiation causes biphasic inflammatory waves in the hippocampus. Also see Supplementary figure 1. **(A)** Experimental design. Sham (SH), n = 6; Irradiation (IR), n = 6, per time point. h = hour, d = day, wk = week, P = postnatal, Hipp = hippocampus. **(B)** Principal component analysis (PCA) plot showing distinct clustering of SH and IR samples from all studied time points. **(C)** Bar plot showing the number of differentially expressed genes (DEGs) between SH and IR animals (*q* < 0.05). Positive y-axis indicates upregulated DEGs; negative y-axis indicates downregulated DEGs. (**D** -**G**) Line graphs showing qPCR analyses of *Ifit1*, *Ifit3*, *Cxcl10* and *Ccl12* in the hippocampus across the studied time points. SH, n = 3-4; IR, n = 3-4. Mean ± SEM, unpaired *t*-test per time per time point to eliminate the animal age factor. *\*p* < 0.05, *\*\*p* < 0.01, *\*\*\*p* < 0.001, *\*\*\*\*p* < 0.0001. (**H** and **I**) Line graphs showing ELISA measurements of CXCL10 and CCL12, respectively, in the hippocampus across the time post-IR. SH, n = 3-5; IR, n = 4-5. Mean ± SEM, unpaired *t*-test per time point. *\*p* < 0.05, *\*\*p* < 0.01.

Gene set enrichment analysis (GSEA) revealed activation (*p* < 0.05) of pathways associated with inflammation and response to viruses (Supplementary figure 1A). Out of the 58 upregulated DEGs, 27 genes were related to inflammation, the majority of which were involved in interferon (IFN) signaling pathways (Supplementary figures 1B and 1C), indicative of a second inflammatory wave mediated by IFN signaling. Targeted expression analysis of genes related to cytokines and chemokines displaying > 3-fold expression changes at any time point revealed that chemokines were the most upregulated inflammatory mediators (Supplementary figure 1D). We found a significant induction of a set of chemokines within the first 24 h post-IR. At 1 wk, *Ccl12* was the only chemokine that remained significantly increased. At 2 wk, however, we detected a significant increase in the expression of *Cxcl10*, *Ccl5* (both belonging to interferon signaling pathways), *Ccl2* and *Ccl12.* The increased expression of *Ccl2* and *Ccl12* remained significant till 6 wk (Supplementary figure 1D). The biphasic inflammatory response was validated in independent hippocampal tissue at both RNA and protein level (Figures 1D-1I; Supplementary figure 1E). These data indicate that WBI causes a biphasic inflammatory response in the hippocampus, where a first wave occurs acutely within the first 24 h post-IR, and a second, delayed wave occurs in the subacute phase, dominated by IFN signaling pathways.

### Delayed microglial responses occurring post-irradiation

We next asked whether the second inflammatory wave detected 2 wk post-IR is driven by a second episode of microglial re-activation. Hippocampal microglia were selectively isolated, based on CX3CR1 expression, at 2 and 6 wk post-IR and processed for bulk RNA-seq ^16^ (Figure 2A). Samples from SH and IR microglia formed clearly distinct clusters at both time points (Figure 2B), indicative of a delayed IR-induced response in microglia. In total, 415 and 1070 genes were differentially expressed at 2 and 6 wk, respectively (Figure 2C; Supplementary list 2), of which only 47 genes were overlapping between the two time points (Figure 2C). GSEA further revealed that microglia from both time points had distinct molecular profiles. At 2 wk, enriched pathways (*q* < 0.05) were related to P53 signaling, cell cycle progression, and response to viral infection (IFN signaling) (Figure 2D); while at 6 wk, enriched pathways were related to cell death (autophagy and necroptosis), innate immune response (TNF signaling, NOD-like receptor signaling, antigen processing and presentation), and response to viral infection (Figure 2E). Targeted analysis of genes related to cytokines and chemokines revealed an increased expression of *Cxcl10* at 2 wk, and *Tnf*, *Il6*, and *Cxcl2* at 6 wk (Supplementary figure 2A).

**Figure 2:**
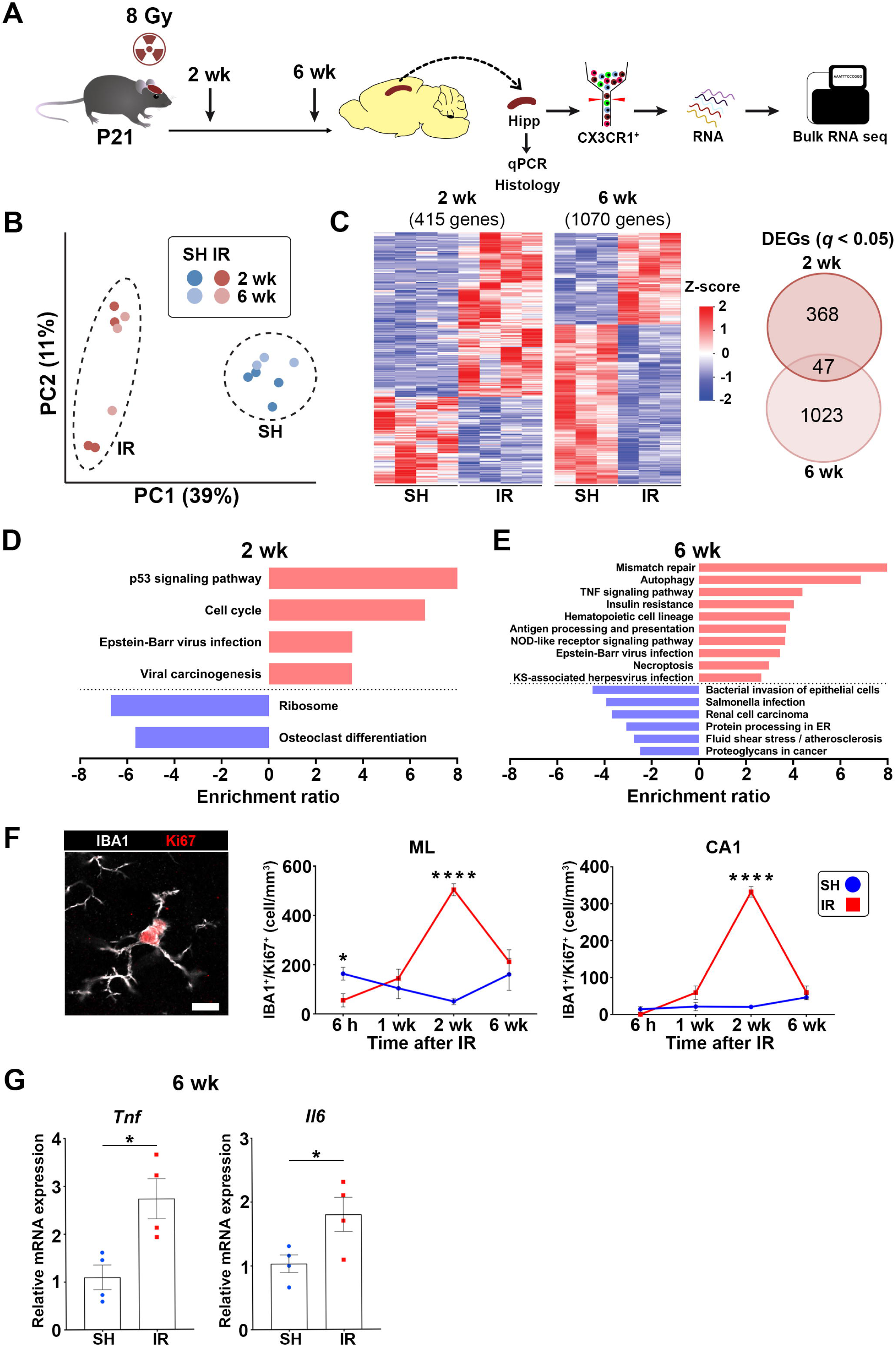
Delayed microglial responses occurring at later phases post-irradiation. Also see Supplementary figure 2. (A) Experimental design. 2 wk, n = 4 per group; 6 wk, n = 3 per group. (B) PCA plot showing distinct clustering of SH and IR samples at both time points. (C) Left: Heatmaps depict DEGs (*q* < 0.05) between SH and IR microglia at 2 and 6 wk. Right: Venn diagram showing the number of distinct and overlapping DEGs between the two time points. (**D** and **E**) Bar plots showing significantly regulated pathways (*q* < 0.05; red = upregulated; blue = downregulated) at 2 and 6 wk, respectively. (F) Left: Representative confocal image displaying co-labeling of microglia (IBA1+, white) with Ki67 (red) in the *cornu ammonis* 1 (CA1) region of the hippocampus 2 wk post-IR. Hoechst (blue), nuclear counter stain. Scale bar = 10 μm. Middle and right panels: Quantification of IBA1^+^/Ki67^+^ cells in the molecular layer (ML) and CA1, respectively, across the studied time points. SH, n = 3-4; IR, n = 3-4. Mean ± SEM, unpaired *t*-test per time point. *\*p* < 0.05, *\*\*\*\*p* < 0.0001.| (G) Bar plots showing qPCR analyses of *Tnf* and *Il6* in the hippocampus 6 wk post-IR. SH, n = 4; IR, n = 4. Mean ± SEM, unpaired *t*-test. *\*p* < 0.05.

To validate these temporal transcriptomic events, we first analyzed the microglial proliferative response post-IR. Proliferating cells (Ki67^+^) were quantified in two distinct hippocampal regions, the molecular layer (ML) and the *cornu ammonis* 1 (CA1) region with the intention to not include the granule cell layer and SGZ, where NSPCs and their progenies often proliferate and microglia actively eliminate apoptotic cells ^29^. The following time points post-IR were covered: 6 h (when the first wave of the inflammation occurred), 1 wk (when the first inflammatory wave was resolved), 2 wk, and 6 wk (during and after the second inflammatory wave). Ki67^+^ cells in these regions were either oligodendrocyte progenitors (OLIG2^+^ cells) or microglia (IBA1^+^ cells) (Supplementary figure 2B). While the total number of Ki67^+^ cells decreased 6 h and 6 wk post-IR (Supplementary figure 2C), phenotyping of Ki67^+^ cells revealed that IR caused a striking shift in the identity of cycling cells in the hippocampus. In the intact hippocampus (SH), OLIG2^+^ cells were the most abundant cycling cells (OLIG2^+^/Ki67^+^), whereas post-IR, cycling cells were predominately IBA1^+^ cells in both examined areas at 2 and 6 wk (Supplementary figures 2D -2E). The number of proliferating microglia (IBA1^+^/Ki67^+^ cells) was significantly increased 2 wk post-IR (Figure 2F), while the number of OLIG2^+^/Ki67^+^ cells decreased over time post-IR (Supplementary figure 2F). Finally, we validated the induction of *Tnf*, *Il6,* and *Cxcl2* 6 wk post-IR in the hippocampus tissue using qPCR (Figure 2G; Supplementary figure 2G). Collectively, these results indicate that microglia undergo a delayed and temporally distinct re-activation post-IR, featured by upregulation of genes involved in IFN signaling, coupled with a wave of proliferation at 2 wk, and adoption of a neurotoxic phenotype at 6 wk.

### Multiple microglial states orchestrate the delayed response to irradiation

Next, we asked whether the observed delayed inflammatory response is exclusively driven by microglia or if other cell types are also involved. To gain better cellular and molecular resolutions, we dissected the hippocampi of IR mice and age-matched SH controls 2 wk post-IR. We improved our previous cell isolation method ^30^ by including two Percoll^®^ gradients for myelin removal to obtain a viable single-cell suspension, containing neurons, suitable for single-cell RNA sequencing (scRNA-seq) using the droplet-based method (10× Genomics Chromium) (Figure 3A). After performing the quality control on the sequenced cells (detailed in the star method section), clustering analysis revealed that our cell isolation protocol captured microglia and macrophages, glial cells (both astrocytes and oligodendrocytes), neurons (both mature and immature), vascular cells (endothelial cells, pericytes, vascular smooth muscle cells (VSMCs), and myofibroblasts), ependymal cells, immune cells, and meningeal fibroblasts, from both SH and IR animals (Supplementary figures 3A - 3D). Microglia were the cells displaying the most DEGs, and with increased expression of inflammatory mediators observed in the hippocampal tissue 2 wk post-IR (*e.g. Cxcl10* and *Ccl12*) (Supplementary figures 3E and 3F). Thus, we focused our attention on analyzing the microglial cells in depth. Microglia from SH and IR were distinctly clustered (Figure 3B). Sub-clustering analysis, considering only the clusters (subpopulations) with significantly enriched proportions (*p* < 0.05) due to the experimental conditions (SH or IR), revealed 12 clusters, of which clusters 1, 3, 4, 6, 7, 9, and 10 were induced post-IR, referred to as radiation-associated microglia (RAM) henceforth (Figure 3C; Supplementary figure 4A; Supplementary list 3). GSEA of signature genes of the RAM clusters revealed activation of pathways related to response to viruses (*i.e.* IFN signaling) in clusters 1, 3, 4, 6, and 10, but more abundant in cluster 6; cell cycle progression (clusters 7, 9, and 10); oxidative phosphorylation (clusters 3, 4, and 9); P53 signaling clusters 1, 6, 9, and 10); cellular senescence (clusters 1, 6, and 10); endocytosis and phagocytosis (cluster 1 and 6) (Figure 3D; Supplementary figure 4B). Representation of activated and suppressed pathways of all RAM clusters, as revealed by GSEA, are shown in (Supplementary figure 5A). These data indicate that RAM are heterogenous and mediate the second inflammatory wave observed 2 wk post-IR.

**Figure 3:**
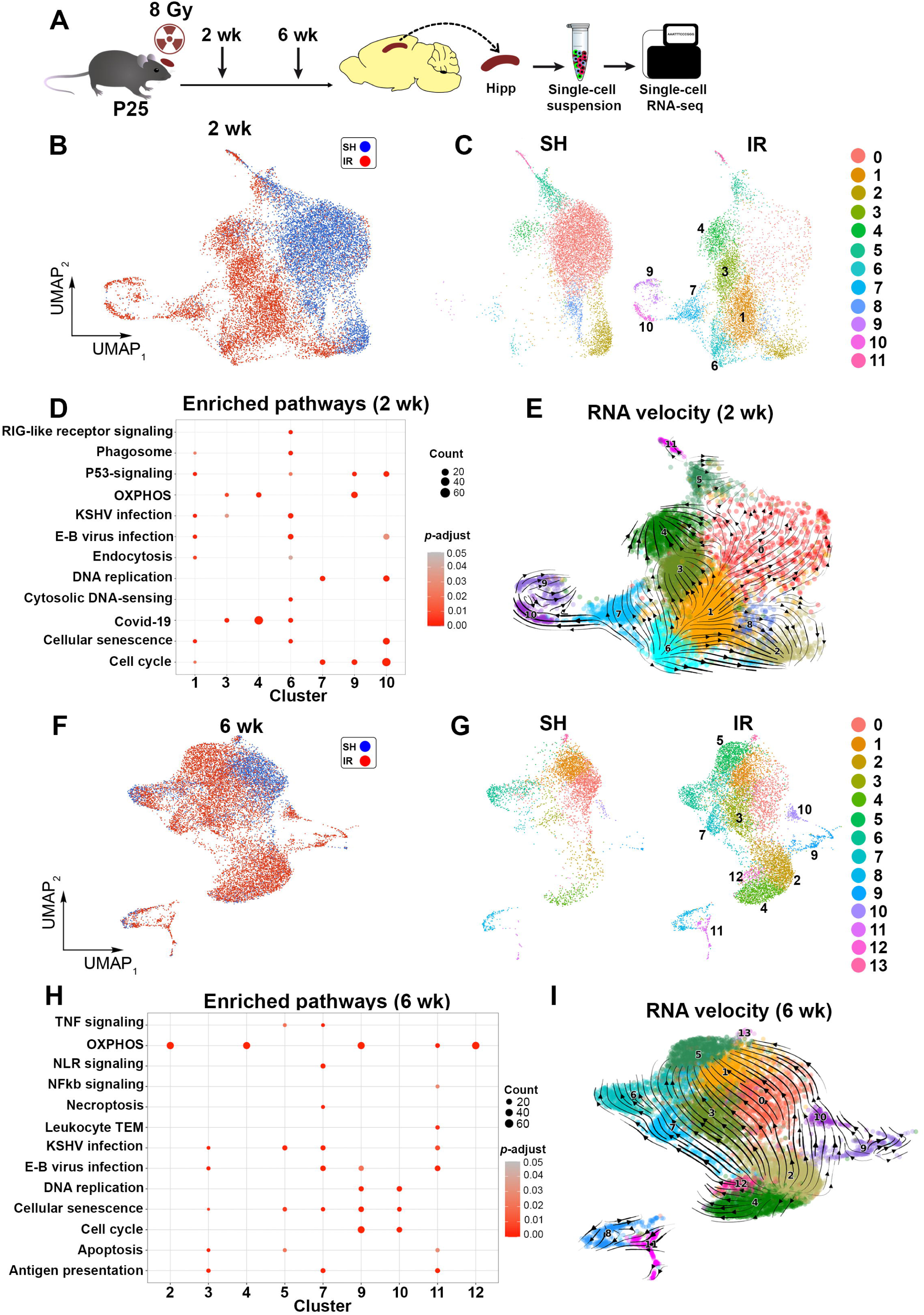
The delayed post-irradiation microglial responses constitute multiple subpopulations. Also see Supplementary figures 3, 4, 5, 6, 7, and 8. **(A)** Experimental design. SH, n = 3; IR, n = 3. **(B)** Uniform manifold approximation and projection (UMAP) showing distinct clustering of hippocampal microglia from SH (blue) and IR (red) 2 wk post-IR. **(C)** Microglial sub-clustering 2 wk post-IR revealed 12 subpopulations. Clusters 1, 3, 4, 6, 7, 9, and 10 were uniquely radiation-associated microglia (RAM) subpopulations. **(D)** Dot plot showing the enriched pathways in RAM clusters (*p* < 0.05) 2 wk post-IR as revealed by GSEA. **(E)** RNA velocity analysis in RAM 2 wk post-IR showing that cells in cluster 6 are drivers for microglial mitotic progression. Arrow lengths indicate the speed of transcriptional activity. **(F)** UMAP showing clustering of hippocampal microglia from SH (blue) and IR (red) 6 wk post-IR. **(G)** Microglial sub-clustering 6 wk post-IR revealed 14 subpopulations. Clusters 2, 3, 4, 5, 7, 9, 10, 11, and 12 represent RAM. **(H)** Dot plot showing the enriched pathways in the RAM clusters (*p* < 0.05) 6 wk post-IR revealed by GSEA. **I)** RNA velocity analysis in RAM 6 wk post-IR showing RAM trajectories towards cluster 5.

We next investigated which RAM subpopulation(s) drive these events. RNA velocity, an algorithm predicting the future states of the individual cells for the coming hours based on the ratio between spliced and non-spliced RNA ^31,32^, revealed that the microglial population with enriched IFN signaling (cluster 6) contained the driver cells of the events seen at this time point, including the mitotic progression (clusters 7, 9, and 10) (Figure 3E).

Next, we looked closer at the microglial population 6 wk post-IR, as they appeared distinctly different from the 2-wk microglia in our bulk RNA-seq data. We applied a similar scRNA-seq approach, and sub-clustering revealed 14 clusters, of which clusters 2, 3, 4, 5, 7, 9, 10, 11, and 12 were RAM (Figures 3F and 3G; Supplementary figure 6A; Supplementary list 3), and the expression of *Tnf* and *Il6*, the two genes that increased post-IR in our bulk data at this time point, was increased in cells belonging to multiple RAM clusters (Supplementary figures 6B and 6C). Again, GSEA of RAM clusters revealed activation of pathways related to IFN signaling and inflammatory responses, cell cycle, cell death, oxidative phosphorylation and leukocyte trans-endothelial migration (TEM) (Figure 3H). Representations of activated and suppressed pathways in all RAM clusters, as revealed by GSEA, are shown in (Supplementary figure 7A). RNA velocity analysis at this time point revealed RAM trajectories towards cluster 5, which was enriched in pathways related to cellular senescence, apoptosis, IFN, and TNF signaling (Figure 3I).

Finally, to relate the molecular events observed in RAM at the 2 and 6 wk time points, we integrated scRNA-seq datasets from these time points. Sub-clustering revealed 10 distinct sub-clusters, with an overlap between the subpopulations detected at both time points; for example, the subpopulations enriched in IFN signaling or cell proliferation. However, their representation was time point dependent, and those uniquely detected at either time point remained distinct. (Supplementary figures 8A - 8C). Together, these results show that specific signaling programs orchestrate the induction of multiple microglial states in the delayed post-IR inflammatory response.

### Activation of the cytosolic nucleic sensing system may contribute to the post-irradiation delayed IFN response

Both genomic instability and mitotic progression with damaged DNA have been shown to result in nucleic acid leakage to the cytosol, generation of micronuclei, and activation of the cyclic GMP-AMP synthase (cGAS) and stimulator of interferon genes (STING) pathway that trigger type-I IFN signaling ^33–35^. We hypothesized that the cGAS-STING pathway may regulate the IFN response observed in microglia 2 wk post-IR due to IR-induced DNA damage and/or mitotic progression with damaged DNA. In a series of proof-of-concept experiments, we leveraged microglial *in vitro* systems where the cells are actively propagating to test this. We first tested whether the presence of micronuclei in microglial cytosol triggers cGAS expression, the upstream component of the pathway. We irradiated microglial cells of different sources (mouse primary microglia, mouse microglial cell line BV2, and the human microglial cell line HMC3) with a single dose of 8 Gy and cultured the cells for at least 24 h before processing for immunofluorescence staining (Figures 4 A and 4B; Supplementary figures 9A - 9C). Next, we used the BV2 and HMC3 cells to perform downstream analyses, since the primary cell cultures yield a limited number of cells and have lower growth potentials. We found that IR significantly increased the fraction of cells expressing the phosphorylated STING (pSTING) in BV2 cells (Figure 4C), and immunoblotting further revealed increased levels of phosphorylated tank-binding kinase 1 (pTBK1), a canonical downstream component of this pathway, in both BV2 and HMC3 cells (Figure 4D; Supplementary figure 9D). Finally, IR resulted in increased expression of the IFN-related genes *Ifit1* and *Ifit3* in both BV2 and HMC3 cells, and *Cxcl10* in BV2, where the levels of the secreted proteins were also increased in the culture media (Figures 4E and 4F; Supplementary figure 9E).

**Figure 4:**
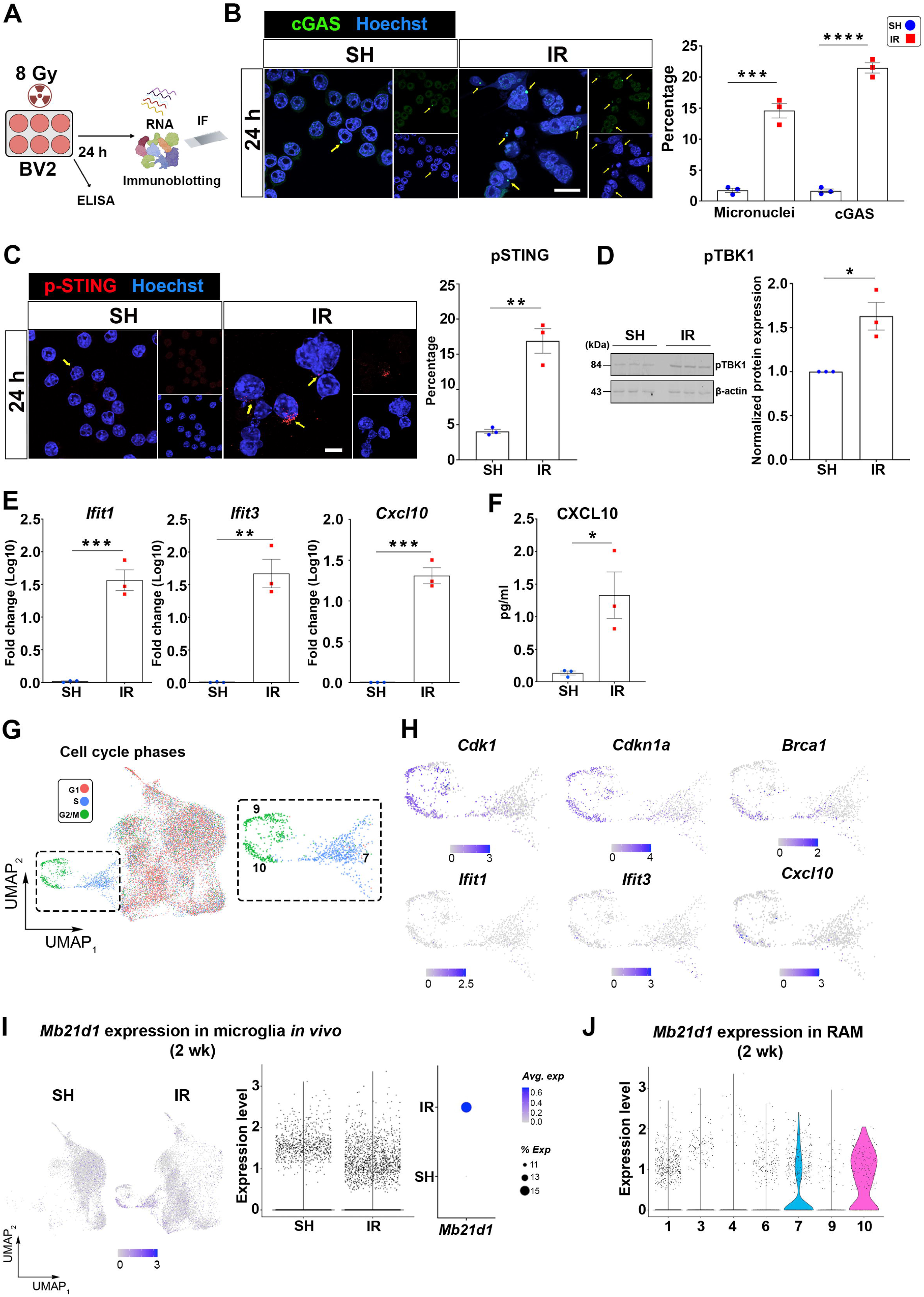
The role of the cytoplasmic DNA sensing on the IFN response post-irradiation. Also see Supplementary figure 9. **(A)** *In vitro* experimental design. IF = immunofluorescence. **(B)** Left: Representative confocal image displaying the micronuclei and expression of cGAS (green; indicated by yellow arrows) in BV2 cells. Hoechst (blue), nuclear counter stain. Scale bar = 20 μm. Right: Bar plots showing the percentage of BV2 cells with micronuclei and cGAS expression 24 h post-IR. Three independent experiments. Mean ± SEM, unpaired *t*-test. *\*\*\*p* < 0.001, *\*\*\*\*p* < 0.0001. **(C)** Left: Representative confocal image displaying expression of phosphorylated STING (pSTING; red; indicated by yellow arrows). Hoechst (blue), nuclear counter stain. Scale bar = 10 μm. Right: Bar plots showing the percentage of BV2 cells expressing pSTING 24 h post-IR. Three independent experiments. Mean ± SEM, unpaired *t*-test. *\*\*p* < 0.01. **(D)** Left: Immunoblots showing the expression of phosphorylated TBK1 (pTBK1) and β- actin expression in BV2 cells in three technical replicates of SH and IR 24 h post-IR from one experimental set. Right: quantification of pTBK1 24 h post-IR from three independent experiments. Mean ± SEM, unpaired *t*-test. *\*p* < 0.05. **(E)** Bar plots showing qPCR analyses of IFN-related genes *Ifit1*, *Ifit3*, and *Cxcl10* in BV2 microglial cells 24 h post-IR. Three independent experiments. Mean ± SEM, unpaired *t*-test. *\*\*p* < 0.01, *\*\*\*p* < 0.001. **(F)** Bar plot showing ELISA measurement of the levels of secreted CXCL10 in the culture media of BV2 microglial cells 24 h post-IR. Three independent experiments. Mean ± SEM, unpaired *t*-test. *\*p* < 0.05. **(G)** Cell cycle analysis of proliferating RAM 2 wk post-IR. **(H)** UMAPs showing expression of genes related to G2/M phase (*Cdk1*), DNA damage response (*Cdkn1a* and *Brca1*), and IFN response (*Ifit1*, *Ifit3*, and *Cxcl10*) in proliferating RAM 2 wk post-IR. **(I)** Left: UMAPs showing expression of *Mb21d1* in microglia from SH and IR animals 2 wk post-IR. Right: Quantification of *Mb21d1*expression in microglia from SH the IR animals 2 wk post-IR. The violin plot shows the expression per individual cell, and the dot plot shows the average expression from all microglia per treatment. **(J)** Violin plot showing quantification of *Mb21d1*expression across the RAM clusters 2 wk post-IR.

*In vitro* IR of the cells causes considerable cell death ^36,37^, which may confound our findings. Hence, we next wanted to validate the results observed in the *in vitro* experiments in our mouse model. We performed cell cycle analyses of proliferating microglial populations in the 2-wk scRNA-seq dataset, since microglial proliferation and IFN-related responses were the major observed events at this time point. We found cells at different cell cycle stages (Figure 4G). Cells in cluster 7 were in S-phase, while those in clusters 9 and 10 had progressed to the G2/M phase (expressing the M phase gene *Cdk1*) despite expressing genes associated with DNA damage response such as *Cdkn1a* and *Brca1*. Subsets of these cells, especially in cluster 10, were expressing the IFN signaling-related genes, such as *Ifit1*, *Ifit3*, and *Cxcl10* (Figure 4H). Moreover, analysis of *Mb21d1* expression (encoding cGAS), as an upstream component of the cytosolic nucleic acid sensing system through all cell types, revealed that it was primarily expressed in microglia and was increased post-IR (Figure 4I; Supplementary Figure 9F). Among RAM, *Mb21d1* expression was detected in several clusters, but most cells with high expression levels were found in the proliferating cells belonging to clusters 7 and 10 (Figures 4I and 4J). These results suggest that microglial mitotic progression while carrying damaged DNA contributes to the delayed IFN response post-IR via activation of the cytosolic nucleic acid sensing system cGAS-STING pathway.

### Monocyte-derived macrophages engraft in the hippocampus and differentiate into microglia-like cells to compensate for the irradiation-induced loss of microglia

Given the dramatic transcriptional profile changes observed in the hippocampal tissue and the microglial cells post-IR, we explored a more chronic phase. Hippocampi were collected 6 months (mo) post-IR, and RNA from the whole tissue was processed for bulk RNA-seq (Figure 5A). We found 30 DEGs between SH and IR groups (*p* adjusted < 0.01), of which 19 were up- and 11 downregulated, respectively. Most downregulated genes were related to microglia, whereas monocyte-derived macrophage (MDM) signatures were upregulated (Figure 5B). To increase the resolution, we performed scRNA-seq on sorted CX3CR1^+^ cells^16^. Clustering analysis revealed 11 clusters, where cluster 1 stood out as distinctly separate from SH and IR microglia (Figure 5C). Cells in this cluster either lacked or had low expression of microglial signature genes, such as *Sall1, Tmem119*, and *Slc2a5*, but expressed MDM-related genes, such as *Itga4*, *Clec12a*, and *Ms4a7* (Figure 5D; Supplementary figure 10A). Notably, to avoid the bias that may result from selectively sorting CX3CR1^+^ cells, we performed scRNA-seq also on unsorted cells, as for the 2 and 6 wk time points, and we confirmed the post-IR MDM recruitment (Supplementary figure 10B). RNA velocity revealed that MDMs display trajectories towards parenchymal microglia, indicating MDM differentiation into a microglial lineage (Figure 5E; Supplementary figure 10C). To validate these results and obtain spatial information on the distribution of MDMs, we performed immunofluorescence staining for IBA1 (labeling both microglia and MDMs) and TMEM119 (marker for parenchymal microglia). We found three distinct populations of IBA1^+^ cells in IR animals, distributed throughout the hippocampus, one population with a high expression of TMEM119 (TMEM119^high^, representing microglia) and two other populations with either negative or lower TMEM119 expression (TMEM119^negative^ and TMEM119^low^, respectively, representing MDMs). IBA1^+^ cells with TMEM119^high^ or TMEM119^negative/low^ were distinctly different, such that TMEM119^negative/low^ cells had thicker processes, less arborization, and smaller area of occupancy (Figure 5F; Supplementary figure 10D). Combined, these data indicate that MDMs engraft the hippocampus long after IR, where they appear to differentiate into microglia-like cells.

**Figure 5:**
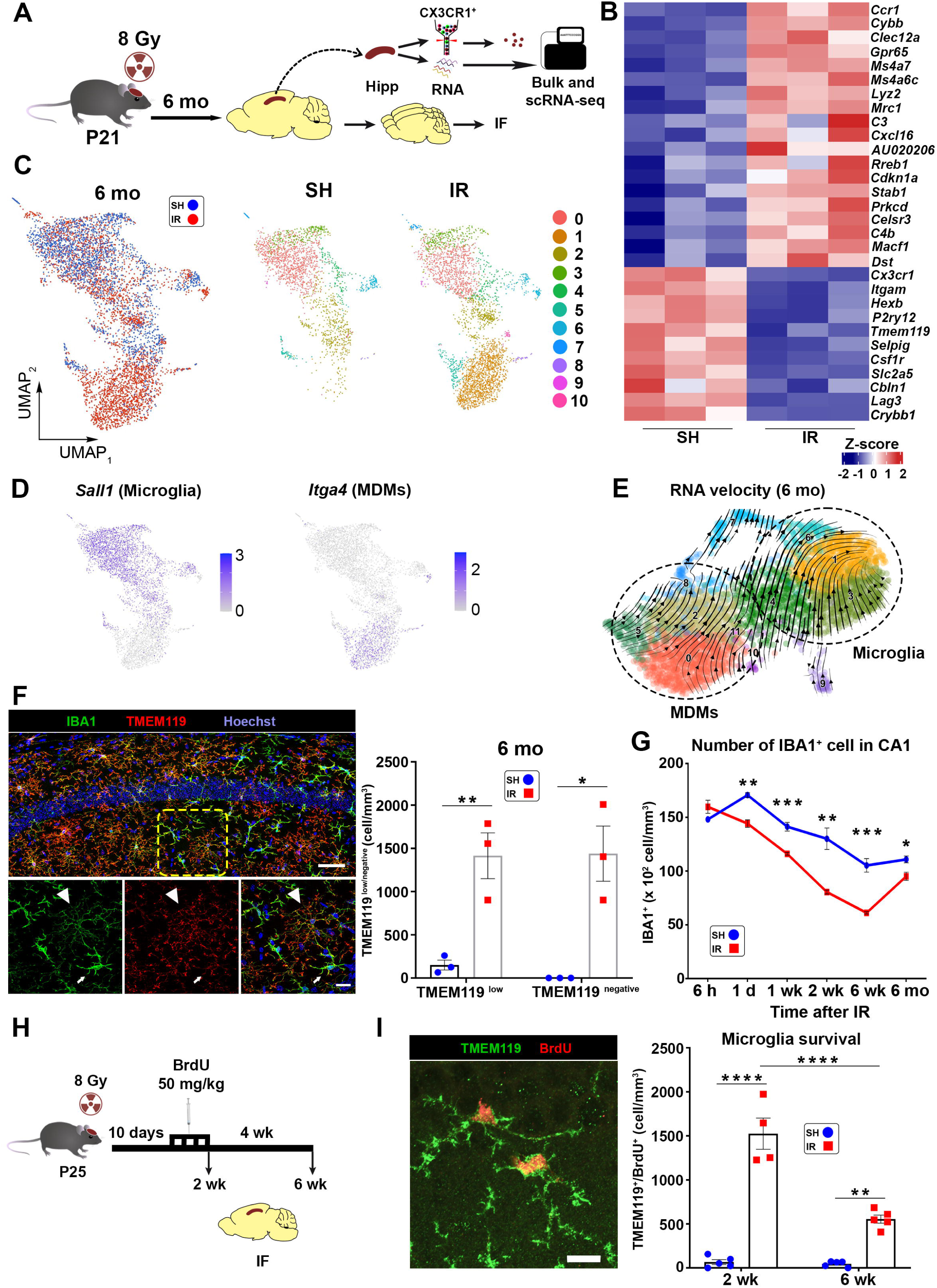
Monocyte-derived macrophages (MDMs) infiltrate the hippocampus to compensate for the microglial loss post-irradiation. Also see Supplementary figures 10 and 11. (A) Experimental design. n = 3 per group. (B) Heatmap depicts DEGs between SH and IR from bulk RNA-seq of the hippocampal tissue 6 mo post-IR. (C) scRNA-seq of isolated CX3CR1^+^ cells 6 mo post-IR Left: UMAP showing clustering of cells from SH (blue) and IR (red). Right: Sub-clustering revealed 11 distinct clusters. RAM in cluster 1 remained distinct from SH and other IR microglia. (D) UMAPs showing expression of signature genes of microglia (*Sall1*) and MDMs (*Itga4*) in RAM 6 mo post-IR. (E) RNA velocity analysis in RAM 6 mo post-IR revealed that MDMs display trajectories towards parenchymal microglia. (F) Left: Representative confocal images displaying co-labeling of IBA1^+^ cells (green) TMEM119^+^ (red) in the CA1 6 mo post-IR. Hoechst (blue), nuclear counter stain. Arrows indicate TMEM119^low^ and arrowheads indicate TMEM119^high^. Scale bar = 50 μm (overview) and = 10 μm (closeup). Right: Bar plot showing quantification of TMEM119^low^ or TMEM119^negative^ cells in the CA1 6 mo post-IR. SH, n = 3; IR, n = 3. Mean ± SEM, unpaired *t*-test. *\*p* < 0.05, *\*\*p* < 0.01. (G) Line graph showing the quantification of IBA1^+^ cells in CA1 region across time post-IR. n = 3-5 per group. Mean ± SEM, unpaired *t*-test per time point. *\*p* < 0.05, *\*\*p* < 0.01, \*\**\*p* < 0.001. (H) Experimental design for assessing microglial survival. (I) Left: Representative confocal image displaying co-labeling of microglia (TMEM119^+^, green) with BrdU (red) in the CA1 2 wk post-IR. Hoechst (blue), nuclear counter stain. Scale bar = 10 μm. Right: Bar plot showing quantification of TMEM119^+^/BrdU^+^ cells 2 wk and 6 wk post-IR in the CA1. 2 wk, n = 4-5; 6 wk, n = 5. Mean ± SEM, two-way ANOVA with Bonferroni’s *posthoc* for multiple comparisons. *\*\*p* < 0.01, *\*\*\*\*p* < 0.0001.

Next, we asked why MDMs infiltrate the hippocampus despite microglia proliferation at earlier time points post-IR. Longitudinal quantification of IBA1^+^ cells post-IR (6 h to 6 mo) demonstrated a progressive loss of IBA1^+^ cells over time, reaching nearly 50% by 6 wk, followed by a near restoration by 6 mo, reducing the loss to 15% (Figure 5G). This led us to hypothesize that IR microglia fail to repopulate through self-renewal, and hence, MDMs infiltrate the hippocampus to compensate for the microglia loss. To test this, we labeled proliferating cells between days 10 and 14 post-IR using 5-bromo-2’-deoxyuridine (BrdU), a thymidine analog incorporated during S-phase ^38^, as these days fall within the proliferative wave shown above. Animals were sacrificed either 2 h after the last BrdU injection (*i.e.* 2 wk post-IR) to assess microglia proliferation, or 4 wk later (*i.e.* 6 wk post-IR) to assess survival of the newborn cells (Figure 5H). Newborn microglia (TMEM119^+^/BrdU^+^ cells) were quantified in the ML and CA1. As expected, IR led to a significant increase in the number of TMEM119^+^/BrdU^+^ cells compared to SH at both time points, but the number of TMEM119^+^/BrdU^+^ cells was reduced by nearly 70% between 2 h and 6 wk post-IR (Figure 5I; Supplementary figure 11A), indicating that the microglia born during the re-activation (*i.e.* 2 wk post-IR) did not survive. Importantly, qPCR analysis of the inflammatory mediators that were increased at earlier time points showed no difference between SH and IR 6 mo post-IR (Supplementary Figure 11B). Together, these results indicate continuous elimination of parenchymal microglia post-IR, failed attempts to compensate for their loss by self-renewal, and consequent repopulation of their empty territories by recruited MDMs that differentiate into microglia-like cells. The observed phenotypic rewiring of the hippocampal resident immune cells post-IR represents a spontaneous elimination of IR-induced neurotoxic microglial phenotypes concurrent with apparent normalization of the inflammatory responses observed at earlier phases post-IR.

### MDMs begin infiltrating the hippocampus early post-irradiation and acquire a microglial phenotype over time

We next computationally investigated the kinetics of the MDM infiltration into the IR hippocampus and their differentiation into microglia-like cells. Analysis of the macrophage populations captured in our scRNA-seq datasets from the 2 wk, 6 wk, and 6 mo post-IR (unsorted cells) identified 8 clusters, of which 3 were IR-induced MDMs (referred to as iMDMs), clusters: 1 (iMDM1), 2 (iMDM2), and 5 (iMDM3) detectable as early as 2 wk post-IR (Supplementary figure 12A). After sub-clustering of iMDMs, iMDM1 yielded two sub- clusters, iMDM1a and iMDM1b (Figures 6A), whose relative abundance varied over time (Figure 6A; Supplementary figure 12B). Using immunofluorescence, we further validated a significant increase in IBA1^+^/TMEM119^negative^ cells in the hippocampal tissue by 6 wk post-IR (Supplementary figure 12C). Next, we performed cell fate mapping using the CellRank- Moscot interface, an algorithm predicting the cellular stages across the experimental timeline based on likelihood ^39,40^. Unsupervised computation for initial states revealed a subset of iMDM3 at 2 and 6 wk as initial iMDMs and subsets of cells belonging to iMDM1 and iMDM2 at 6 mo as terminal states (Figure 6B; Supplementary figure 12D). To confirm these findings, we assessed the expression of the microglia signature gene *Tmem119* in iMDMs and found minimal expression in iMDM3 (the initial state of iMDMs), intermediate in iMDM1b, and higher in iMDM1a and iMDM2, indicating that iMDM1 and iMDM2 were indeed the iMDMs that had differentiated into microglia-like cells (Figure 6C). Moreover, morphological analysis of the IBA1^+^/TMEM119^negative^ and IBA1^+^/TMEM119^low^ cells revealed increased arborization of these cells at 6 mo compared to 6 wk (Supplementary figure 12E), compatible with the acquisition of a more microglia-like morphology over time.

**Figure 6.**
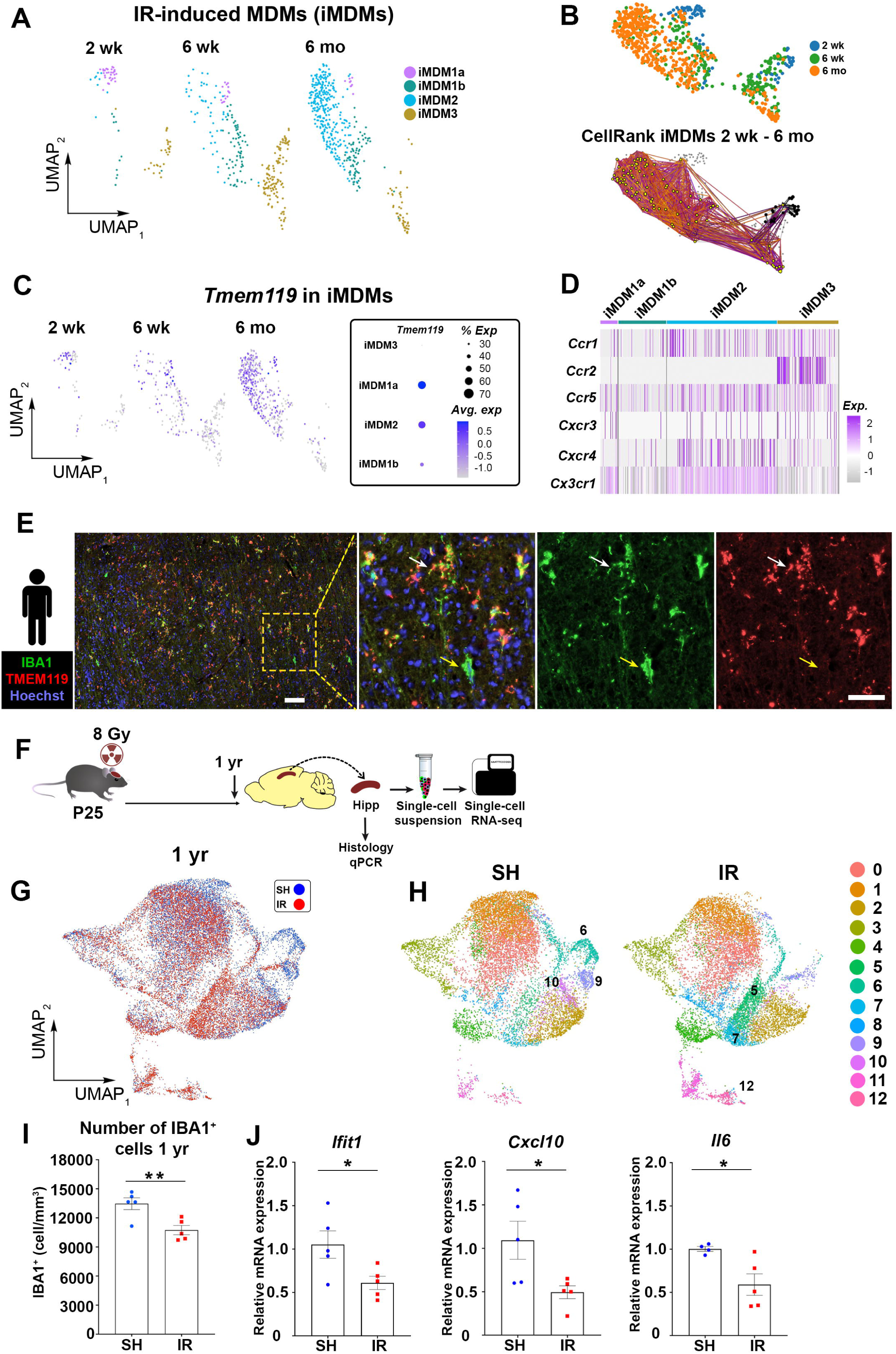
: MDMs infiltrating the brain at an early phase post-irradiation. Also see Supplementary figures 12, 13, 14, and 15. (A) UMAP showing the dynamics of IR-induced MDM (iMDM) clusters over time post-IR. (B) Top: UMAP combining iMDM clusters from 2 wk, 6 wk, and 6 mo. Bottom: CellRank- Moscot analysis revealed that iMDMs from 2 and 6 wk (black dots) are the initial state, and cells in 6 mo (yellow dots) are the terminal state. (C) Left: UMAPs showing *Tmem119* expression in iMDMs over time post-IR. Right: Dot plot showing the magnitude of *Tmem119* expression in the iMDM clusters. (D) Heatmap showing the expression of chemokine receptors in the iMDM clusters. (E) Representative images displaying the expression of IBA1 (green) TMEM119 (red) in cancer-free hippocampus of a medulloblastoma patient 5 years post-radiotherapy. Yellow arrows indicate IBA1^+^/TMEM119negative cells; white arrows indicate IBA1^+^/TMEM119^+^ cells. Hoechst (blue), nuclear counter stain. Scale bar = 100 μm (overview) and = 50 μm (closeup). (F) Experimental design. yr = year. n = 3 per group. (G) UMAP showing clustering of hippocampal microglia from SH (blue) and IR (red) 1 yr post-IR. (H) UMAPs showing sub-clustering of hippocampal microglia 1 yr post-IR. Thirteen clusters were revealed, of which some were enriched (clusters 5 and 7) and some repressed (cluster 6) post-IR. (I) Bar plot showing quantification of IBA1^+^ cells in the CA1, 1 yr post-IR. SH, n = 5; IR, n = 5. Mean ± SEM, unpaired *t*-test per time point. *\*\*p* < 0.01. (J) Bar plots showing qPCR analyses of inflammatory mediators *Ifit1*, *Cxcl10*, and *Il6* in the hippocampus 1 yr post-IR. SH, n = 4-5; IR, n = 5. Mean ± SEM, unpaired *t*-test. *\*p* < 0.05.

Next, we performed an unsupervised analysis of chemokine receptor expression in iMDMs as a proxy for possible signaling axes for recruiting these cells to the hippocampus post-IR. We found that *Ccr2* was expressed by initial state cells (iMDM3) but not by iMDM1 and 2 (Figure 6D). iMDM3 displayed minimal expression of *Cx3cr1*, a microglia signature gene whose expression was higher in iMDM1 and iMDM2 (Figure 6D), further supporting iMDM3 being the initial state and in line with a previous report ^41^ demonstrating a role for CCR2 in recruiting MDMs into the brain. We also found a subset of iMDM3 expressing *Ccr1, Ccr5*, *Cxcr3*, and *Cxcr4* (Figure 6D). This suggests a role for these chemokine receptors in MDM recruitment, in addition to CCR2, supported by significantly increased IBA1^+^/TMEM119^negative^ cells 6 wk post-IR also in mice lacking *Ccr2* (Supplementary figure 12F). *Ccr1*, *Ccr5*, and *Cxcr4* expression remained in a subset of iMDM2 (differentiated into microglia-like cells) (Figure 6D).

Next, we wanted to investigate the source of iMDMs. Recent work has demonstrated both blood and CNS-associated bone marrow in the skull and vertebrae as reservoirs for CNS infiltrating myeloid cells under pathological conditions ^42^. Thus, we compared the iMDMs from the 6 wk time point, containing both initial *Ccr2*-expressing and differentiated cells, with the previously published scRNA-seq data of CNS-infiltrated MDMs recruited from blood or CNS-associated bone marrow ^42^. We found that iMDMs overlapped with subsets in both populations, suggesting that both sources serve as the origin of iMDMs (Supplementary figures 13A -13C).

Finally, in cancer-free hippocampal tissue sections from an autopsied brain collected 5 years after craniospinal irradiation of a medulloblastoma patient, we observed two distinct microglial populations, IBA1^+^/TMEM119^+^ and IBA1^+^/TMEM119^negative^ (Figure 6E; Supplementary figures 14A -14C). This observation suggests that infiltration of MDMs post-IR also occurs in humans, but further validation is required. Collectively, our data indicate that MDMs infiltrate the hippocampus early post-IR and differentiate into microglia-like cells over time.

### Microglial alterations post-irradiation persist long-term, but without active inflammatory responses

We used scRNA-seq to explore the molecular profiles of microglia at a much later time point, one year (yr) post-IR (Figure 6F). SH and IR microglia overlapped to a greater extent compared with all earlier time points, with some exceptions (Figure 6G). Sub-clustering revealed 13 clusters, of which clusters 6 and 10 were significantly enriched in SH animals, and cluster 6 was nearly absent in IR animals (Figures 6G and 6H; Supplementary figure 15A). GSEA of cluster 6 revealed enrichment of pathways related to neurons and synapses (Supplementary figure 15B). Significant RAM clusters were 5, 7, and 12 (Figures 6G and 6H). Clusters 5 and 7 were enriched in pathways related to oxidative phosphorylation, protein processing, synapses, and lysosomes, whereas cluster 12 was enriched in MDM genes (Supplementary figures 15B and 15C). Although cluster 9 did not statistically differ between SH and IR, the clustering pattern in SH and IR animals differed. At this late time point, the number of IBA1^+^ cells was still significantly lower (∼15%) post-IR (Figure 6I), and IR animals tended to have lower expression of genes encoding inflammatory mediators, significantly so for *Ifit1*, *Cxcl10*, and *Il6* (Figure 6J; Supplementary figures 15D and 15E). Together, these data demonstrate that the post-IR microglial loss persists, and long-term alterations of microglia appear not related to the production of inflammatory mediators, but rather to phagocytosis and synaptic regulation.

## The second wave of microglial responses appears crucial for the long-term effects of irradiation on microglia

To connect the temporal molecular events occurring in RAM, we integrated our scRNA-seq data from unsorted microglia obtained from the 2-wk, 6-wk, 6-mo, and 1-yr time points, both RAM and age-matched SH. The integrated cells generated 9 distinct clusters, clustering based on their molecular similarities across the time points (Supplementary figures 16A and 16 B; Supplementary list 3). Molecular trajectories can be masked by integration of datasets, so to better visualize these across the mouse lifespan, we instead favored merging the datasets, and this generated 12 distinct clusters (Figures 7A and 7B; Supplementary figure 16C; Supplementary list 3). SH microglia clustered in an age-dependent manner (Figure 7A). For RAM, however, the developmental patterns were perturbed, and subsets of the 2-wk, 6-wk, 6- mo, and 1-yr RAM clustered together (Figure 7A), suggesting that the alterations of RAM were induced during early phases post-IR.

**Figure 7:**
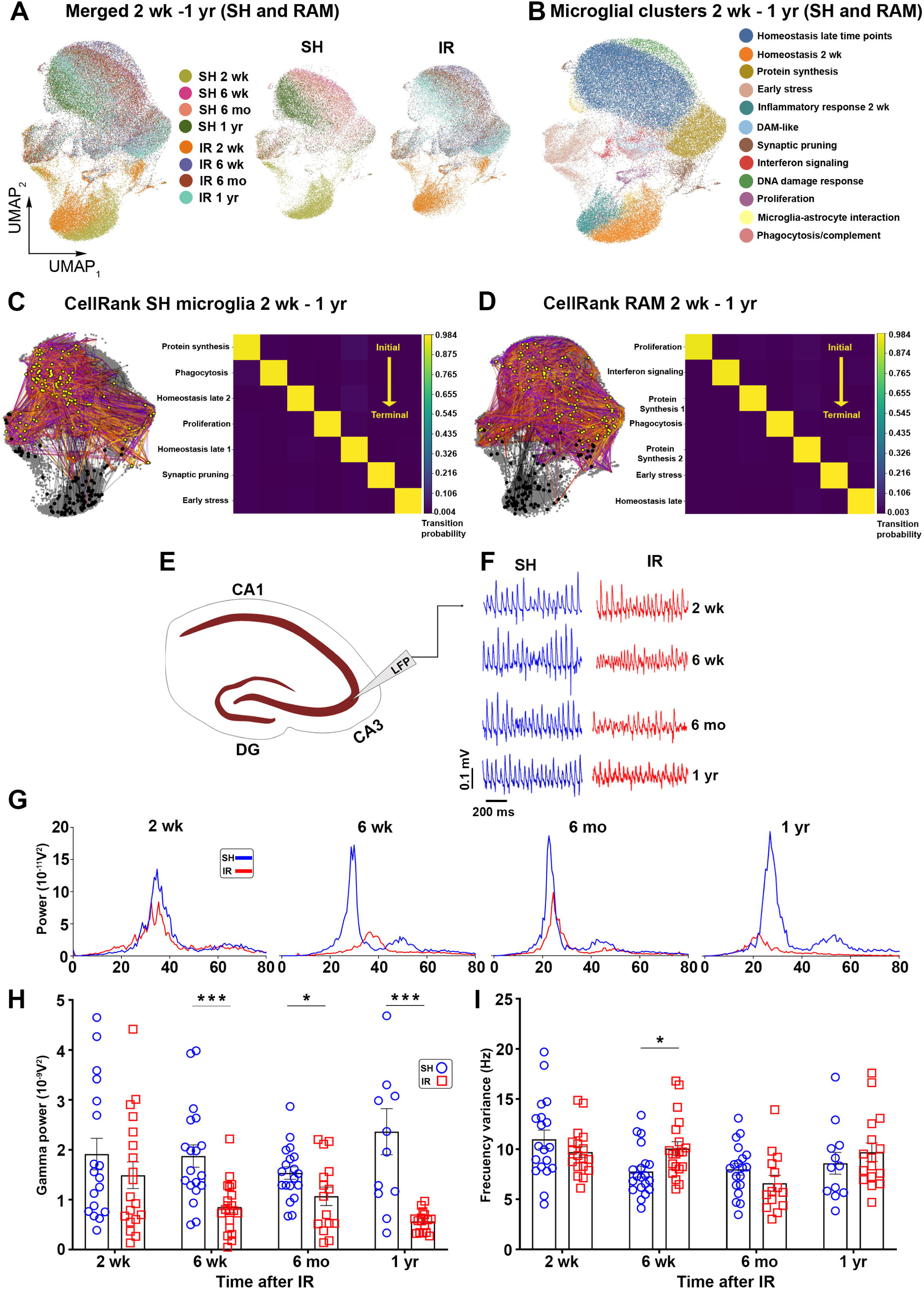
Longitudinal overview of RAM (2 wk – 1 yr) and the hippocampal neuronal network synchrony. Also see Supplementary figure 16. (A) UMAPs showing merging of RAM and their age-matched SH control microglia from the 2-wk to 1-yr time points. (B) Sub-clustering of the merged microglia from the 2-wk to 1-yr time points. Clusters were annotated based on the DEGs. (C) CellRank-Moscot analysis of SH microglia over time. Left: Visualization of the random walk of the initial (black dots) and the terminal (yellow dots) states. Right: Coarse-grained transition matrix showing the probabilities of initial and terminal states. (D) CellRank-Moscot analysis of RAM over the time post-IR. Left: Visualization of the random walk of the initial (black dots) and the terminal (yellow dots) states. Right: Coarse- grained transition matrix showing the probabilities of initial and terminal states. (E) Representative diagram of electrode location in area CA3 of hippocampal slices. (F) Representative sample traces of local field potentials (LFP) recordings from SH and IR slices 2[wk, 6[wk, 6[mo, and 1 yr. SH, n = 3-5; IR, n = 3-5, per time point. (G) Representative power spectra of hippocampal CA3 network activity recorded in SH and IR slices 2[wk, 6[wk, 6[mo, and 1 yr. (H) Gamma oscillation power post-IR. Mean ± SEM, unpaired *t*-test per time point to eliminate the animal age factor. \**p* < 0.05, \*\**p* < 0.01, \*\*\**p* < 0.001. (I) Frequency variance at the above-mentioned time points after IR. Mean ± SEM, unpaired t- test. \**p* < 0.05.

We reconstructed the molecular trajectories in SH microglia and RAM over time using the CellRank-Moscot tool by setting the 2-wk microglia as the initial state. In SH animals, terminal states were confined populations within the 1-yr time point, and intermediate trajectories between 6-wk and 6-mo time points were observed before committing to a terminal state (Figure 7C; Supplementary figure 16D). In RAM, the terminal states were scattered between several clusters identified in cells belonging to the 6-mo and 1-yr time points, and the intermediate trajectories also appeared dispersed (Figure 7D; Supplementary figure 16D). Importantly, the proliferating RAM and those enriched in genes related to IFN signaling were determined as initial states (Figure 7D). These results suggest that the uncovered, delayed IFN signaling and mitotic progression are crucial for the long-term effects of IR on hippocampal function.

### Hippocampal circuitry asynchrony occurs 6 wk and 1 yr post-irradiation

Lastly, we wanted to correlate the microglial rewiring and inflammatory dynamics to neuronal function. To this end, we evaluated hippocampal gamma oscillatory activity 2 wk, 6 wk, 6 mo, and 1 yr post-IR (Figures 7E and 7F), allowing for evaluation of neuronal firing patterns and circuitry rhythmicity involved in cognitive behaviors ^43,44^. We found that the power and rhythmicity (as judged by the frequency variance) in the gamma range were comparable between the IR and the SH animals 2 wk post-IR (Figures 7F-7I). At 6 wk, however, we detected a profound decrease in gamma oscillation power in IR hippocampi, concurrent with increased frequency variance (Figures 7F-7I). At 6 mo, the gamma oscillation power had partially recovered in IR hippocampi (Figures 7F-7I), and differences in rhythm regularity (*i.e.* frequency variance) were normalized at this time point (Figures 7F- 7I). Interestingly, we found a subsequent robust decrease in the gamma oscillation power 1 yr post-IR (Figures 7F-7I).

## Discussion

Microglia shape the physio-anatomical neuronal properties through synaptic pruning and secretion of trophic factors to maintain proper neuronal function ^14,45–47^. Perturbed microglial-neuronal interaction due to microglial hyperreactivity or dysfunction may cause neuronal dysfunction, reflected in impaired cognition, as documented in numerous CNS disease models ^14^ . Impaired cognition is a hallmark complication in CNS cancer survivors treated with cranial radiotherapy ^1^. Microglia play pivotal roles in IR-induced neurotoxicity through the secretion of inflammatory molecules, generation of reactive oxygen species, and excessive synaptic elimination ^18^. Despite this consensus in the field, there is no successful treatment targeting microglial responses to prevent or reverse IR-induced cognitive deficits. Here, we uncover numerous hippocampal RAM states induced over time post-IR. We show that RAM are highly heterogeneous and follow unique temporally-regulated molecular programs resulting in multifaceted inflammatory profiles. IFN signaling has been implicated in the development of cognitive deficits in various CNS disease models ^48–50^. In our IR model, we observed biphasic inflammatory waves in the hippocampal tissue, an acute wave occurring within hours post-IR and a delayed wave after 2 wk, featuring IFN signaling concurrent with microglial mitotic progression. This second peak is followed by increased levels of genes encoding the neurotoxic cytokines TNFα and IL-6 6 wk post-IR, coinciding with neuronal network asynchrony, a combination of events often linked to impaired cognitive performance in CNS disease models ^51–53^. We previously showed that hippocampal circuits are compromised in mouse models of neurodegenerative diseases, where microglia play pivotal roles ^43,54,55^, and targeting neuroinflammation rescues the hippocampal gamma oscillatory activity and cognitive performance in these models ^56^. The neuronal asynchrony was partially recovered at 6 mo, overlapping with spontaneous elimination of irradiated parenchymal microglia, repopulation with MDMs, and normalized levels of genes encoding TNFα and IL-6 in the hippocampal tissue. Microglia depletion before IR did not affect hippocampal neurogenesis ^57^, a process important for memory and learning, but elimination of microglia post-IR has been shown to yield improved cognitive performance in animal models when assessed shortly after repopulation ^26,58^. However, the duration of this improvement remains unknown, especially considering the subsequent neuronal asynchrony observed 1 yr post-IR without increased expression of inflammatory mediators, rather the opposite. This strongly argues against a role for chronic inflammation late post-IR, making it unlikely that the second neuronal asynchrony is linked to active inflammatory responses. Featured regulated pathways at this time point were related to phagocytosis and neuronal synapses, indicating synaptic dysregulation by microglia. Nevertheless, the delayed neuronal asynchrony may still be a secondary consequence of the initial inflammatory events post-IR, leading to engagement of other cell type(s) mediating these late effects, as was previously shown in the context of chemotherapy-induced neurotoxicity ^59^. Adverse effects of IR to the brain are not limited to microglia or inflammation; other consequences, such as impaired neurogenesis, demyelination, gliosis, vascular abnormalities, and direct damage to neurons, are expected to contribute ^1^.

Radiation causes DNA damage and genomic instability, and cells that fail to repair their DNA may undergo cell death or cellular senescence ^60,61^. Senescent cells often resist cell death, do not replicate, and secrete soluble factors, known as senescence-associated secretory phenotype (SASP), affecting the functions of neighboring cells ^62^. Microglia express the DNA damage response and cell cycle arrest genes *Cdkn1a* and *Ccng1* early post-IR ^16^, but despite this, previous work ^12^ and this study demonstrated increased microglial proliferation. Mitotic progression with damaged DNA or chromosomal instability activates the cGAS-STING pathway, leading to the induction of type-I IFN signaling ^33,63^, resulting in cellular senescence in neighboring cells via the production of SASP ^64,65^. In this study, we uncovered subsets of microglia characterized by IFN signaling and mitotic progression concurrent with damaged DNA. Our *in vitro* experiments showed that IR can activate the cytosolic DNA sensing system in microglia, as judged by increased expression of cGAS, pSTING, and pTBK1. *In vivo*, we found that the observed IFN response 2 wk post-IR preceded the increased levels of *Tnf* and *Il6* 6 wk post-IR, both classified as SASP factors ^66^. Thus, future studies targeting the components of the cGAS-STING pathway to dampen the post-IR IFN response are warranted.

Our results demonstrated rewiring of hippocampal parenchymal phagocytes after IR. The recruitment of MDMs into the brain post-IR has been reported earlier using histological and bulk RNA-seq, where MDMs were shown to infiltrate the brain post-IR and differentiate into microglia-like cells ^67–70^. Here, we provide single-cell resolution and longitudinal characterization of the kinetics and the molecular phenotypic transition into microglia-like cells. Our results not only confirm the role of CCR2 as a key chemokine receptor for recruiting MDMs into the postnatal brain ^41^, but also suggest a role for other chemokine receptors, such as CCR1 and CCR5, in this process. The detection of MDMs in the hippocampus of *Ccr2-*deficient mice post-IR further supports this, is consistent with what is found in non-CNS tissue ^71^, and also explains the earlier finding that increased *Ccr1* expression post-IR remained significantly higher in *Ccr2-*deficient mice ^72^. These chemokine receptors may synergize to achieve efficient MDM recruitment into the hippocampus post-IR. Indeed, we observed elevations of CCL2 and CCL12, both of which are ligands for CCR1 and CCR2. While CCL2 followed the biphasic inflammatory pattern, CCL12 remained significantly elevated at the protein level for at least 6 wk post-IR, when the initially recruited cells (iMDM3) were abundant, suggesting a key role for CCL12 in MDM recruitment to the hippocampus post-IR. Comparison of the molecular profiles of the post-IR MDMs with previously published datasets of blood- or CNS-associated bone marrow-derived MDMs ^42^ suggested that both sources are the origin of MDMs. Previous work has demonstrated blood-derived MDM infiltration into the IR brain by systemic injection of bone marrow-derived cells ^70^. A cell tracing approach would be necessary to conclusively establish the CNS-associated bone marrow as a reservoir of MDMs post-IR.

We provide a single-cell level characterization of microglial responses from 2 wk to 1 yr after cranial IR. The data serve as a resource for defining novel biomarkers associated with brain IR, measured in cerebrospinal fluid or blood. The uncovered re-activation of microglia 2 wk post-IR, characterized by IFN signaling and proliferation despite DNA damage, offers potential pathways to counteract subsequent negative consequences on the hippocampal and brain functions. However, given the anti-tumor properties mediated by the IFN signaling ^73^, such approaches should be considered with caution. Finally, given that MDMs and parenchymal microglia display functional differences in animal models mimicking bacterial infection ^67,68^, it would be relevant to investigate how the rewired phagocytes react to a secondary brain injury, such as re-irradiation or tumor relapse.

## Supporting information

Supplementary list 1

Supplementary list 2

Supplementary list 3

## Acknowledgments

We express our deep gratitude to the patients and families who donated clinical samples, and to the Swedish Childhood Tumor Biobank (Barntumörbanken) for providing patient materials. KB was supported by the Swedish Childhood Cancer Fund (Barncancerfonden), the Swedish Cancer Foundation (Cancerfonden), the Swedish Research Council, grants provided by the Stockholm Region (ALF projects), Radiumhemmets Forskningsfonder, the Frimurare Barnhus Foundation in Stockholm, the Karolinska Institute Doctoral (KID) funding, the KI Foundation for Research, the Märta and Gunnar V. Philipson Foundation, and the Swedish Brain Foundation (Hjärnfonden). AO was supported by Stiftelsen Samariten, the KI Foundation for Research, and Erik Rönnbergs Stipendium. VML was supported by the Swedish Research Council [grant numbers: 2021-02801 and 2023-03015] and the Robert Bosch Stiftung (Stuttgart, Germany). CB was supported by the Karolinska Institutet, Center for Innovative Medicine grant (CIMED). AF was supported by Stiftelsen Samariten, KI Foundation for Research, Mary Béves Stiftelse för Barncancerforskning, ìShizu Matsumuraîs Donation. EP was supported by Stiftelsen Barnforskningen Astrid Lindgrens Barnsjukhus.

The authors acknowledge support from the National Genomics Infrastructure in Stockholm funded by Science for Life Laboratory, the Knut and Alice Wallenberg Foundation, and the Swedish Research Council. We thank SNIC/Uppsala Multidisciplinary Center for Advanced Computational Science for assistance with massively parallel sequencing and access to the UPPMAX computational infrastructure.

We are grateful to Hajar Ba Omar, Yiqun Tang, Mercedes Posada Pérez, Marie-Kim St-Pierre, João Pedro Alves-Lopes, and Jan-Bernd Stukenborg for sharing their expertise on troubleshooting immunofluorescence staining of the patient and postmortem tissues. We would also like to thank Guillermo Vázquez Cabrera for helping to set up the HMC3 human microglia cells culture. We would also like to thank the following core facilities at the Karolinska Institutet: the X-ray Irradiation Core Facility, the Flow Cytometry Core Facility for assisting with the microglia sorting, the Bioinformatics and Expression Analysis (BEA) for bulk RNA-seq; the Eukaryotic Single Cell Genomics Facility (ESCG) for running the scRNA-seq. Illustrations were partly created using BioRender.com.

## Author contributions

A.M.O. and K.B. conceived and designed the study. A.L.R., E.P., A.M.O. A.Fr., and K.B. interpreted the results and wrote the manuscript. A.L.R., E.P., G.A.Z., K.Z., C.F.D.R., and A.M.O. performed the animal experiments. A.L.R., G.P., G.A.Z., T.S., and A.M.O. performed the histological analyses. A.L.R., E.P., and A.M.O. performed the isolation of microglia and whole-brain cells for transcriptomic analyses. A.L.R., T.S., and Y.R. performed the qPCR.

K.Z. Y.X., and C.Z. contributed to bulk RNA-seq data. A.L.R., N.O-V., and V.M.L. analyzed the bulk RNA-seq data. E.P. performed single-cell RNA-seq and computational analyses. Y.S. and C.B. contributed to the single-cell RNA-seq analysis. G.P., L.E.A.G., A.G.I., and A.Fi. performed and analyzed the neuronal gamma oscillations. A.L.R., Y.R., T.S., M.Q.C., L.F., and B.J. contributed to the BV2, HMC3, and immunoblotting experiments. L-M.C., A.S., and K.B. reviewed and annotated the clinical data. All authors discussed the results, commented on, or edited the manuscript.

## Disclosure of Potential Conflicts of Interest

V.M.L. is co-founder and owner of HepaPredict AB. The other authors declare no competing of interests.

## Materials and methods

### Experimental model and subject details Human samples

The clinical samples used in this study were provided by the Swedish Children’s Tumor Bank (Barntumörbanken; ethical application nr. 2021-02333). The cancer-free hippocampal tissue was obtained from the autopsied brain tissue of a 12-year-old patient with Group 4 medulloblastoma collected 5 years post-radiotherapy. The tumor-bearing tissue was collected at surgical resection of an ependymoma from a 4-year-old patient upon a tumor relapse 1 year and 10 months after the initial surgery. The healthy human hippocampal tissue used as a control for the immunofluorescence staining was purchased from Clinisciences (Biochain; #T2234052). This tissue was donated by a healthy 54-year-old male. BioChain’s tissue products are based on the sample repository network established following the approved ethical standards and procedures. All tissues were paraffin-embedded and cut into 5 μm-thick sections.

### Animals

Female mice were used in all studies, as irradiation (IR)-induced cognitive deficits are more severe in females, both in animal models and patients ^2,74^. Twenty-one- or twenty-five-day-old mice of the following strains were used C57BL/6J (Charles River, Sulzfeld, Germany, stock #000664), B6.129(Cg)-*Ccr2*tm2.1Ifc/J (*Ccr2*^RFP^) (Jackson laboratory; stock #017586). Animals were housed in equal light/dark cycles (12/12 h) and were fed *ad libitum*. All the experimental procedures were carried out according to the European and Swedish animal welfare regulations approved by the northern Stockholm ethical committee (application nr. N248/13, N141-16 and 13676-2020).

### Irradiation procedure

Mice were initially anesthetized with 5% isoflurane in an induction chamber in a mixture of air and oxygen (1:1), then transferred to the irradiation machine and placed in a prone position. The anesthesia was maintained with 1.5 % isoflurane during the irradiation procedure. The following irradiation machines were used: X-RAD 320 (PXi Precision X-ray, North Branford, CT, USA) or CIX3 cabinet X-ray irradiator (Xstrahl, Surrey, England). For the X-RAD 320 irradiator, the animal head was distanced approximately 50 cm from the radiation source, and an irradiation field of 2 × 2 cm was used to cover the entire head. Animals received a single dose of 8 Gy delivered at a rate of 0.73 Gy/min. For the Xstrahl irradiator, the animal head was irradiated with a circular field of 1.5 cm in diameter. The collimator was an Xstrahl Perspex tip applicator. A single dose of 8 Gy was delivered with a dose rate of 1.349 Gy/min (dosimetry uncertainty ∼ 2%) at 300 kV and 10 mA. Focus skin distance (FSD) to the animal head was 32 cm. External filtration giving a half-value layer (HVL) of 4.0738 mm Cu was applied by adding a Thoraeus filter (1.0 mm Sn, 0.25 mm Cu, 1.50 mm Al). Litter-mates sham controls (SH) were subjected to the same duration of anesthesia in the absence of IR. Animals were allowed to recover from the anesthesia and returned to their cages.

### 5-bromo-2’-deoxyuridine (BrdU) administration

To assess the survival of proliferating microglia, dividing cells were labeled between day 10 and 13 post-IR with the thymidine analog BrdU (SigmaAldrich #B5002). BrdU was injected i.p. at a dose of 50 mg/kg, two injections per day (6 h apart). Animals were sacrificed either 2 h or 4 weeks after the last BrdU injection to assess the proliferation and survival of the newborn cells, respectively.

### Tissue collection

Animals were sacrificed at 6 hours, 1 day, 1 week, 2 weeks, 6 weeks, 6 months, or 1 year post-IR. Animals were deeply anesthetized with sodium pentobarbital (100 mg/kg, ABCUR AB #444362, Sweden) and transcardially perfused with 1× phosphate-buffered saline (PBS; ThermoFisher Scientific #10010023). Brains were collected and the hemispheres were separated. The left hemispheres were placed into 4% paraformaldehyde (PFA; Histolab Products # HL96753.1000, Sweden) and stored at 4°C for 48 h. The brains were then cryoprotected in 30% sucrose solution (Sigma-Aldrich #S7903) made in 0.1 M phosphate buffer pH 7.4 and stored at 4°C until slicing. Right hemisphere hippocampi were dissected and kept in a -80 °C freezer until further processing for RNA or protein extraction.

### Generation of single-cell suspension for RNA sequencing

Single-cell suspensions were prepared from three SH controls and three IR mice per time point. Animals were deeply anesthetized with sodium pentobarbital (100 mg/kg, ABCUR AB, Sweden) and transcardially perfused with ice-cold 1× PBS without Ca^2+^and Mg^2+^(PBS; pH 7.4; Gibco/Life Technologies #10010056). Brains were collected and the hippocampi from the two hemispheres were dissected and put into 1.5 mL eppendorf tubes containing 1× PBS placed on ice. The tissue was chopped into small pieces using scalpel and transferred into conical tubes containing an enzymatic mixture of Dispase II (0.01%; Sigma-Aldrich #D4693), papain (0.1%; Roche #000000010108014001), and DNaseI (0.05%; Roche # 000000010104159001) in 1× Hank’s buffered salt solution (HBSS) without Ca^2+^and Mg^2+^ (Gibco/Life Technologies #14175095), later supplemented with 12.4 mM magnesium sulfate (Sigma-Aldrich #M7506). Tubes were incubated at 37°C for 10 min, after which the enzymatic activities were stopped with 20% ice-cold heat-inactivated fetal bovine serum (FBS; Gibco/Life Technologies #10500064). The cell suspension was then filtered through a 70 μm cell strainer and centrifuged for 5 min at 500 × g at 4°C. After washing with 1× HBSS, cells were resuspended in 20% percoll solution (percoll plus, GE Healthcare, #GE17-0891- 02); 10× phenol red HBSS (Gibco/Life Technologies #14060040) and 1× HBSS, and overlaid with an equal volume of 1× HBSS. Tubes were spun at 1,000 × g at 4°C for 30 min with no break to remove myelin. The cell pellet was resuspended in 1 ml flow cytometry buffer (R&D Systems #FC001). In protocols intended to collect neurons, an additional centrifugation step in a higher percoll concentration was applied. The supernatant from the first percoll step (20%) was collected in 50 ml conical tubes and mixed with an equal volume of 50% percoll solution to obtain a concentration of 30 % percoll, and spun at 1,000 × g at 4°C for 30 min with a break. The supernatant was removed, and the pellet was resuspended in 1 ml flow cytometry buffer. Both cellular fractions collected from the two-step percoll were mixed and spun at 500 × g at 4°C for 5 min. The pellet was resuspended in 0.5 ml flow cytometry buffer. Cells processed for single-cell RNA-seq were pooled from three SH and three IR animals before adding the flow cytometry buffer and processed for 10× single-cell RNA sequencing. For the sorted microglia, only the first percoll step was applied to get the cell suspension that contained the microglial fraction. That cell suspension was incubated with a Fc-blocker (Miltenyi # 130-092-575) for 10 min, followed by incubation with a PE-conjugated goat anti-CX3CR1 antibody (R&D Systems #FAB5825P) for 15 min. PE-conjugated goat IgG (R&D Systems #IC108P) was used as isotype negative control. SytoxRed (Invitrogen #S11381) was used as a live/dead cell stain. Cells were sorted using FACSAria III (BD Biosciences, New Jersey, USA) into a 1.5 ml eppendorf tube containing 0.5 ml flow cytometry buffer and placed on ice, when processed for 10×single-cell RNA-seq, or a 1.5 mL eppendorf tube containing 0.25 ml Qiazol (Qiagen #79306) and placed on dry ice, then stored -80°C. For each time point, cells were isolated from SH and IR animals at the same time and processed in the same manner (as one batch).

## Cell culture

For the primary hippocampal microglia culture, fourteen-day-old mouse pups were deeply anesthetized with sodium pentobarbital (100 mg/kg, ABCUR AB, Sweden) and transcardially perfused with ice-cold 1× PBS without Ca^2+^and Mg^2+^(PBS; pH 7.4; Gibco/Life Technologies #10010056). Brains were collected, and the hippocampi from the two hemispheres were dissected and put into 1.5 mL eppendorf tubes containing 1× PBS placed on ice. The tissue was chopped into small pieces using a scalpel and transferred into conical tubes containing an enzymatic mixture of Dispase II (0.01%; Sigma-Aldrich #D4693), papain (0.1%; Roche #000000010108014001), and DNaseI (0.05%; Roche # 000000010104159001) in 1× HBSS without Ca^2+^and Mg^2+^ (Gibco/Life Technologies #14175095), later supplemented with 12.4 mM magnesium sulfate (Sigma-Aldrich #M7506). Tubes were incubated at 37°C for 10 min, after which the enzymatic activities were stopped with 20% ice-cold heat-inactivated fetal bovine serum (FBS; Gibco/Life Technologies #10500064). The cell suspension was then filtered through a 70 μm cell strainer and centrifuged for 5 min at 500 × g at 4°C. After washing with 1× HBSS, cells were resuspended in 20% percoll solution (percoll plus, GE Healthcare, #GE17-0891-02); 10× phenol red HBSS (Gibco/Life Technologies #14060040) and 1× HBSS, and overlaid with an equal volume of 1× HBSS. Tubes were spun at 1,000 × g at 4°C for 30 min with no break to remove myelin. Cells were seeded in T25 cell culture flasks in DMEM/F12 with Glutamax culture medium (Gibco/Life Technologies #31331028) containing 10% heat-inactivated FBS and supplemented with 10 ng/ml recombinant mouse M-CSF (R&D systems 416-ML-010) and grown at 37°C in 5% CO_2_. The culture medium was changed every second day. After reaching approximately 90% confluence, cells were washed with 1× PBS and incubated with 0.05% trypsin-EDTA (Gibco/Life Technologies) for 5 min at 37°C. The enzyme activity was stopped with FBS, and cells were washed with 1× PBS. Cells were resuspended in flow cytometry buffer (R&D Systems #FC001) and processed for magnetic separation using CD11b micro-beads (Miltenyi Biotec #130-093-636) following the manufacturer’s recommendations. Cells were washed and seeded in 12-well plates at a density of approximately 2 × 10^4^ cultures in the above-mentioned cell culture medium, however, without M-CSF and left to settle for 48 h prior to irradiation.

For the microglial cell lines, the murine microglia BV2 and human HMC3 cell lines were used. BV2 cells were cultured in DMEM GlutaMAX (GIBCO/Life Technologies #10564011). HMC3 cells were cultured using EMEM media with L-glutamine (ATCC #30- 2003). Cells were supplemented with 10% FBS and 5% penicillin/streptomycin, grown at 37°C in 5% CO_2_, split and passaged every 2 - 3 days. For irradiation, BV2 cells were seeded in 6-well plates at a density of 1.5 × 10^5^ and allowed to settle overnight. HMC3 cells were seeded either in 6-well plates, 1 × 10^5^ cells per well in 3 ml of media (for RNA collection), or in T-25 flasks with 2.5 × 10^5^ cells per flask (for protein extraction), and allowed to settle overnight. Cells were irradiated in a CIX2 X-ray irradiator cabinet (Xstrahl, Surrey, England) with a single dose of 8 Gy delivered at a rate of 1.35 Gy/min at 195 kV and 10 mA. Focus skin distance (FSD) to the flask /plate was 40 cm. External filtration giving an HVL of 9.0436 mm was applied by adding a 3.0 mm Al filter. A rotating platform was used to ensure homogenous dose delivery. Control cells were positioned in the rotating platform for a similar amount of time required to deliver 8 Gy without IR. Irradiated cells and their respective sham controls were grown for 1 h (for BV2), 24 h (for BV2 and primary culture), or 48 h (for HMC3) post-IR. The culture media were collected and spun for 10 min at 4°C at 10,000 × g, and the supernatants were transferred into 1.5 ml tubes and stored at -80°C. The cells were washed with 1× PBS, and the following buffers were added depending on the intended downstream analysis: protein extraction buffer (for western blot), RLT buffer containing 1% β-mercaptoethanol (for RNA expression) or ice-cold 4% PFA (for immunostaining) and processed for the intended downstream analysis.

## RNA isolation and quantitative real-time polymerase chain reaction (qPCR)

RNA was isolated from the cultured cells using the RNeasy Plus Micro Kit (Qiagen, #74034; for BV2) and RNeasy Plus Mini Kit, Qiagen, #74136; for HMC3). For samples obtained from the *in vivo* experiments, RNA was extracted using the RNeasy Plus Mini Kit (Qiagen, #74134; from the hippocampal tissue), and the miRNeasy Micro Kit (Qiagen, #217084; from the sorted cells). cDNA was generated using the QuantiTect® Reverse Transcription Kit (Qiagen, #205311) using the AB Simpliamp Thermal PCR machine (Applied Biosystems). The qPCR was performed using the QuantiTect SYBR Green PCR Kit (Qiagen, #204143). Target mRNA expression was assessed using the QuantiTect primer assay (Qiagen). The following primers were used: Mm_*Gapdh*_3_SG (#QT01658692); Mm_*Ifit1*_1_SG (#QT01161286); Mm_*Ifit3*_1_SG (#QT00292159); Mm_*Cxcl10*_1_SG (#QT00093436); Mm_*Ccl12*_1_SG (#QT00244391); Mm_*Tnf*_1_SG (QT00104006); Mm_*Il6*_1_SG (#QT00098875); Mm_*Cxcl2*_1_SG (QT0011325); Hs_*GAPDH*_1_SG (#QT00079247); Hs_*IFIT1*_1_SG (#QT00201012); Hs_*IFIT3*_1_SG (#QT00100030);_ Hs_*CXCL10*_1_SG (#QT01003065). qPCR was performed using either the StepOnePlus™ Real-Time PCR System (Applied Biosystems #4376600) or the CFX384 Touch Real-Time PCR Detection System (Bio-rad #1855484). Relative mRNA expression was determined using the ^ΔΔ^CT method, and *Gapdh* was used as a housekeeping gene.

## Bulk RNA sequencing

Isolated RNA was processed for sequencing at the Bioinformatics and Expression Analysis core facility (BEA) at Karolinska Institutet, Sweden, or at BGI Genomics, China. For sequencing of RNA isolated from the whole hippocampus tissue, the total RNA was subjected to quality control with Agilent Tapestation according to the manufacturer’s instructions. 200 ng of total RNA were subjected to Illumina sequencing, and libraries were prepared with the Illumina TruSeq Stranded mRNA ligation prep kit, which includes cDNA synthesis, ligation of adapters, and amplification of indexed libraries. The yield and quality of the amplified libraries were analyzed using Qubit by Thermo Fisher and the Agilent Tapestation. The indexed cDNA libraries were normalized and combined, and the pools were sequenced on the Illumina Nextseq 550 using a V2.5 75-cycle flowcell generating 75 bp single-end reads. Basecalling and demultiplexing were performed using BCL2 software with default settings generating Fastq files for further downstream mapping and analysis. For sequencing of RNA isolated from sorted microglia at 2 and 6 wk post-IR, the RNA quality was checked on an Agilent bioanalyzer 2100 using the Eukaryote total RNA picochips, and library preparation was performed using QIAseq FX single-cell RNA library kit (QIAGEN), then sequenced on an Illumina HiSeq 2000. For data analysis, unsupervised hierarchical clustering and principal component analyses (PCA) were performed in Qlucore Omics Explorer 3.2 (Qlucore, Lund, Sweden). Differentially expressed genes were determined by comparing irradiated- and sham-control animals using heteroscedastic two-tailed *t*-tests. Multiple testing correction was performed using the Benjamini-Hochberg algorithm with a false discovery rate (FDR) of 5%. For targeted expression analysis of genes related to cytokines and chemokines, genes displaying > 3-fold expression changes at any time point were included.

Data from the whole hippocampus tissue obtained 6 months post-IR were analyzed using R (version 4.3.0). The DESeq2 (version 1.40.2) package was used to identify differentially expressed genes across different groups. Log fold-changes obtained with a Wald test were shrunk using the apeglm method. Genes with less than 10 counts were excluded. Differentially expressed genes were identified by considering adjusted *p* < 0.1 as calculated with the Benjamini-Hochberg multiple adjustment method.

For gene set enrichment analysis, ranked gene lists were prepared based on signal to noise ((Mean irradiated - mean sham control)/sum of standard deviations). Human gene names were utilized for input of ranked gene lists into the GSEA application https://www.gsea-msigdb.org/gsea/ index.jsp, Broad Institute, Inc), using the GSEAPreranked tool.

## Single-cell RNA sequencing

Isolated cells were processed for single-cell RNA sequencing at the eukaryotic single-cell genomics facility (ESCG) at Karolinska Institutet, Sweden, using the 10× Genomics Chromium method (version 3) and sequenced using Illumina NovaSeq 6000 with an S1, S2 or S4 flow cell. The Cellranger 6.0.1 pipeline was used to align the raw sequencing reads to the mouse reference genome, *Mus musculus* version *mm10,* and generate the unique molecular identifier (UMI) count matrix. The generated count matrix was analyzed using the Seurat R package (version 3.0.0). Each condition included 3 technical replicates, and each replicate was processed independently before integration. Low-quality cells were excluded based on the number of genes expressed in the cells and the percentage of mitochondrial counts. Cells expressing a minimum of 250 genes and a maximum of 6,000 genes were considered for downstream analysis, given that each gene was expressed in at least three cells. The hippocampal cellular components are heterogeneous, and each cell type has a different energy demand, with neurons having the highest energy consumption ^75^. To ensure inclusion of all cell types, no threshold for the mitochondrial counts was included during the first cleanup. The filtered count matrices were normalized by dividing the feature counts for each cell by the total number of counts from all cells, then multiplied by 10,000 and log-transformed. Next, we identified highly variable features for each dataset independently. We used the Canonical Correlation Analysis (CCA) - default Seurat method for the integration of datasets and removal of the batch effect. To ensure optimal results, we separately integrated our datasets using Harmony integration. Both methods yielded similar results, so we proceeded with CCA. In brief, we used canonical correlation analysis and identified “anchors” (conserved cell groups) between datasets, which were used for batch correction and the comparison of the differentially expressed genes between experimental conditions. The data were scaled using Pearson residuals, regressing out the variation by the mitochondrial genes. PCA was then performed on the scaled data for the dimensionality reduction, using 30 components. The output was used to produce a Uniform Manifold Approximation and Projection for Dimension Reduction (UMAP) with a resolution of 0.5 for identification of the clusters. Identification of each cluster was based on signature genes of each cell type according to the existing literature as well as through the automatic cell type recognition package SingleR. Signature genes for microglia included *Sall1*, *Cx3cr1*, *P2ry12, Tmem119*; for macrophages *Mrc1* and *Ms4a7*; for oligodendrocytes, *Pdgfra*, *Mbp, Olig2,* and *Mog*; for astrocytes *Gja1, Gfap,* and *Aqp4*; for neurons *Dcx*, *Rbfox3, Neurod1,* and *Syt1*; for endothelial cells *Cldn5*; for pericytes *Vtn* and *Pdgfrb*; for fibroblasts/myofibroblasts *Col1a1, Acta2, and Lama1*; for B cells *Cd79a, Cd19 and Igkc*; for T- and natural killer cells *Cd7, Cd3g,* and *Itgb7*. The proliferating cells were identified using markers such as *Mki67* and *Top2a*. At this stage, we revisited the levels of the mitochondria and customized them for each cell type separately. Specifically, microglia threshold was set to 10 %, astrocytes/radial glia to 40%, and oligodendrocytes, ependymal, pericytes, endothelial, VSMCs/fibroblasts, B/T/NK cells, meningeal fibroblasts to 20%. After the second quality control, a total of 124,879 single cells collected from the 2-week, 6-week, 6- month, and 1-year time points were analyzed. Cell numbers per condition per time point per experimental condition for the two-week all cell types: SH: 25,969; IR: 25,903. Two-week microglia: SH: 8,149; IR: 7,988. Six-week microglia: SH: 4,107; IR: 10,099. Six-month microglia (CX3CR1 sorted): SH: 2,191; IR: 3,533. Six months microglia (unsorted): SH: 5,940; IR: 8,190. One-year microglia: SH 11,304; IR: 11,506. As we focused on microglia, the microglial cells from all studied time points were selected, sub-clustered and analyzed in- depth.

The Speckle package was used to analyze differences in cell type proportions across experimental conditions and time points. Using the *propeller* function, we looked for significantly different clusters between SH and IR groups, based on ANOVA statistical testing.

ClusterProfiler package was used for gene set enrichment analysis of the upregulated and statistically significant differentially expressed genes (DEGs) of the microglial clusters from each time point. Using the compareCluster function, we directly compared the enriched functional profiles of each cluster using the KEGG database. DEGs were taken into account when expressed by at least 30% of cells in the cluster, the log2 fold-change was higher than 0,8, and the *p*-adjusted value was lower than 0.05.

Scoring of the cell cycle phases of each cell was calculated using the CellCycleScoring function in the Seurat pipeline, based on a set of genes expressed during the S and G2/M phases. The assigned scores for the S and G2/M phases were saved on the metadata of the Seurat object.

For the estimation of the RNA velocity, the post-alignment bam files were used to run velocyto command v0.17, *run* for any technique. To mask the potential repeats of expressed genes, we retrieved the mouse repeat annotation file from the UCSC genome browser (mm10_rmsk.gtf). The mouse genome annotation reference was acquired from CellRanger, and the resultant loom files contained the spliced and unspliced counts. The scVelo tool was used to estimate the generalized RNA velocity model. We extracted the coordinates, count matrix, and gene names of the processed Seurat object for microglia from R and created an anndata object. The obtained loom file and the anndata object were merged, and the generalized dynamical model was used for the velocity estimation and plotting of velocities. The Cellrank-Moscot module was used to examine the development of the cells across experimental time points, since RNA velocity is incapable of such computing. According to the default procedure of Moscot to perform time series data analysis, we set up a temporal problem, which matched cells coming from different time points using the method of optimal transport (OT)^40^. A kernel (RealTimeKernel) was created to compute the transition matrix using a connectivities’ self-transition weight of 0.2. Setting the 2-week timepoint as the initial state, we inferred transition probabilities and cellular dynamics, which we visualized through random walks.

We passed the RealTimeKernel to the Generalized Perron Cluster Cluster analysis (GPCCA) estimator to estimate and classify macrostates into initial, intermediate, and terminal states based on the real Schur decomposition and computed 7 states in the SH or IR groups, respectively. We plotted the relationships between the predicted macrostates in a coarse-grained transition matrix, ranked based on the stability probability of each state. We then predicted terminal states (g2.predict_terminal_states), using a stability threshold of 0.97 and proceeded to compute fate probabilities (g2.compute_fate_probabilities) towards terminal states, which we plotted onto the original UMAP of SH or IR groups.

## Immunohistochemistry and immunofluorescence

For mouse brain tissues, the left hemispheres were cut sagittally into 25-μm-thick free-floating sections made in 1:12 series intervals using a sliding microtome (Leica SM2010R) and stored in 2 ml eppendorf tubes containing a cryoprotectant solution (25% glycerol, 25% ethylene glycol in 0.1M phosphate buffer) and kept at +4°C. After several washes with 1× Tris-buffered saline (TBS), samples were incubated in sodium citrate solution (NaCi, 10 mM, pH 6.0) for 30 min at 80°C when antigen retrieval was needed. When immunoperoxidase staining was used, sections were incubated in a 0.6% hydrogen peroxide (H_2_O_2_) solution for 30 min to quench the endogenous peroxidase. For BrdU staining, the double-stranded DNA was denatured by incubating the sections in 2 N HCl at 37°C for 30 min followed by a neutralization step performed by incubating the section in a 0.1 M borate buffer for 10 min at room temperature. Sections were washed with TBS. Non-specific binding was blocked by incubating the sections in a solution of 3% or 5% normal donkey serum (Jackson ImmunoResearch Laboratories; #017000121), 0.1% Triton X-100 (made in TBS) for 1 h at room temperature. Sections were then incubated with primary antibodies at 4°C for 24 - 72 h, depending on the antibody. The following primary antibodies were used: goat anti-IBA1 (Abcam# ab5076; 1:500); rabbit anti-IBA1 (Wako Chemicals #1919741; 1:1,000); rabbit anti-TMEM119 (Abcam #ab209064; 1:500); rat anti-Ki67 (ThermoFisherScientific #14- 5698-82; 1:500); goat anti-OLIG2 (R&D Systems #AF2418; 1:500); rat anti-BrdU (Abcam #6326; 1:500). Sections were incubated for 1 h or 2 h at room temperature with appropriate biotinylated or fluorescent secondary antibodies, respectively. The following secondary antibodies were used: Biotinylated donkey anti-goat IgG (Jackson ImmunoResearch #705- 065-147; 1:1,000); AlexaFlour-488 donkey anti-goat IgG (Molecular probes/Life Technologies #A11055; 1:1,000); AlexaFlour-488 donkey anti-mouse IgG (Molecular probes/Life Technologies #A21202; 1:1,000); AlexaFlour-555 donkey anti-rabbit IgG (Molecular probes/Life Technologies #A31572; 1:1,000); AlexaFlour-555 donkey anti-rat IgG (Abcam #ab150154; 1:1,000); CF-633 donkey anti-goat IgG (Biotium #20127; 1:1,000). To visualize the immunoperoxidase staining, sections were incubated for 1 h at room temperature in avidin-biotin solution (Vectastain ABC Elite kit, Vector Laboratories #PK6100, Burlingame, CA; 1:100). The color precipitate was developed with a solution containing H_2_O_2_, nickel chloride, and 3-3’diaminobenzidine tetrahydrochloride (DAB; 1:100; Saveen Werner AB, Malmö, Sweden). Sections were mounted into slides, dehydrated with NeoClear (Merck #1.09843.5000, Germany) and coverslipped using NeoMount mounting medium (Merck #1.09016.0500, Germany).

For human tissues, sections were deparaffinized using xylene, rehydrated in descending ethanol series (100 - 70%), and then washed with 1× TBS. Antigen retrieval was performed by heating the section in a solution containing 10 mM Trisma base (Sigma-Aldrich), 1 mM ethylenediaminetetraacetic acid (EDTA; Merck-Millipore) and 0.05% tween-20 (Merck- Millipore), pH 9. Non-specific binding was blocked by incubating the sections in a 1× TBS solution containing 5% normal donkey serum (Jackson ImmunoResearch Laboratories, #017000121), 0.1% Triton X-100 (Sigma-Aldrich; #X100) for 1 h at room temperature. Sections were incubated with goat anti-IBA1 (Abcam #ab5076; 1:500) and rabbit anti- TMEM119 (Abcam #Ab185333; 1:500) overnight at 4°C. After several washes with 1× TBS, sections were incubated for 2 h at room temperature with the following secondary antibodies: AlexaFluor-488 donkey anti-goat IgG (Molecular probes/Life Technologies #A11055; 1:1,000) and AlexaFluor-555 donkey anti-rabbit IgG (Molecular probes/Life Technologies #A31572; 1:1,000).

For cultured cells, after fixation (explained above) and several washes with 1× TBS, non- specific binding was blocked by incubating the sections in a solution of 5% normal donkey serum (Jackson ImmunoResearch Laboratories), 0.1% Triton X-100 (made in 1× TBS) for 1 h at room temperature. Cells were incubated overnight with the following primary antibodies: mouse anti-γH2AX (Phospho S139; Abcam #Ab26350; 1:500); rabbit anti-cGAS (D3O8O; Cell signaling technology #31659; 1:250); rabbit anti-cGAS (E5V3W; Cell signaling technology #79978; 1:250); rabbit anti-phospho-STING (Ser365; Cell signaling technology #62912; 1:250). Cells were incubated for 1 h at room temperature with the appropriate secondary antibody. The following antibodies were used: AlexaFluor-488 donkey anti-rabbit IgG (Molecular probes/Life Technologies #A-21206; 1:1,000); AlexaFluor-555 donkey anti-rabbit IgG (Molecular probes/Life Technologies #A31572; 1:1,000); CF-555 donkey anti-mouse IgG (Biotium #20037; 1:1,000).

Hoechst 33342 (Molecular Probes/Life Technologies #H3570) was used as a nuclear counterstain. Slides were coverslipped using ProLong Gold anti-fade reagent (Molecular probes/Life Technologies; #P36930).

## Microscopy and cell quantification

All histological analyses were performed in the molecular layer (ML) and the *Cornu Ammonis* 1 (CA1) region of the hippocampus in sections containing the dorsal hippocampus spaced 300 μm apart (*i.e.* every 1:12 series). Analyses of total Ki67 positive (Ki67^+^) cells, proliferating microglia (IBA1^+^/Ki67^+^) or Oligodendrocyte progenitors (OLIG2^+^/Ki67^+^), parenchymal microglia (IBA1^+^/TMEM119^high^) and infiltrated macrophages (IBA1^+^/TMEM119^negative^ or IBA1^+^/TMEM119^low^) were performed using the LSM 700 Zeiss confocal scanning microscopy (Carl Zeiss, Germany), equipped with the Zen software (Black edition 2012, Carl Zeiss). Z-stack images were acquired in sequential scans performed at 1 μm section intervals using a 20× objective lens and 1 airy pinhole setting and analyzed using Zen Blue Lite software (Carl Zeiss; Germany). The fluorescent intensities of TMEM119^+^ cells observed in the microglia of sham controls were considered TMEM119^high^, and signals below those levels were considered TMEM119^low^. The total numbers IBA1^+^ cells over the time post-IR, were quantified using a bright field microscope (AxioImager M2; Carl Zeiss, Germany) equipped with the StereoInvestigator software (MicroBrightField Inc.), or in images were acquired using ZEISS Axio Scan.Z1 slide scanner (Carl Zeiss, Germany). In all quantifications, the total number of cells was the sum of all counted cells in all sections per animal multiplied by the series interval (*i.e.* 1:12). The cell density was determined by dividing the total number of quantified cells by counting volume.

Morphological analyses of parenchymal microglia (IBA1^+^/TMEM119^high^) and microglia-like cells (IBA1^+^/TMEM119^negative^ ^or^ ^low^) were performed on the Imaris software (Imaris V9.6). Confocal z-stack images were processed for 3-D reconstruction, and individual IBA-1 positive cells were isolated using the surface tool. The filament tracer tool was then applied to every masked surface. The mean diameter of the processes, area coverage, and number of branching points were then analyzed.

For analysis of micronuclei and cGAS expression in SH or IR cultured microglial cells, images were acquired using the LSM 700 laser scanning confocal microscope (Carl Zeiss, Germany) equipped with the Zen software (Black edition 2012, Carl Zeiss) for primary cells and BV2, and LSM 900 laser scanning confocal microscope (Carl Zeiss, Germany) equipped with the Zen blue software (Carl Zeiss, Germany) for HMC3. For analysis of phospho-STING, images were acquired using the ZEISS Axio Scan.Z1 slide scanner (Carl Zeiss, Germany). For each SH or IR condition, three coverslips were analyzed. Per coverslip, at least seven random fields were imaged covering the borders and the center of the coverslip. Image analysis was performed using the Zen Blue Lite software (Carl Zeiss, Germany). To analyze the percentage of cells with concomitant micronuclei and cGAS expression, at least 2,300 and 1,100 cells (Hoechst^+^) from SH and IR cells were analyzed, respectively. A micronucleus is defined as a discrete Hoechst^+^ DNA aggregate apart from the primary nucleus of the cell. For analysis of phospho-STING expression, at least 1,200 and 700 cells from SH and IR cells were analyzed, respectively.

## Protein extraction, ELISA, and immunoblotting

For protein extraction from the hippocampal tissues, ice-cold protein extraction buffer (50 mM Tris-HCl; Sigma-Aldrich #T1503, 100 mM NaCl; Sigma-Aldrich, #S7653; 5 mM EDTA, Sigma-Aldrich #E5134 and 1 mM EGTA, Sigma-Aldrich #E3889) supplemented with protease inhibitor cocktail (Roche #11836170001) and phosphatase inhibitors (Roche #04906837001) were added to the frozen tissue and homogenized with a sonicator. Samples were then centrifuged for 10 min at 4°C at 10,000 × g, and the supernatants were transferred into 0.5 ml tubes and stored at -80°C. The total protein concentration was determined using the Pierce BCA protein assay kit (Thermo Fischer Scientific #23225), and absorbance was measured using the FLUOstar Omega (BMG LABTECH, Germany) plate reader.

The levels of the chemokines in the hippocampal homogenates were measured using the following ELISA kits: mouse IP-10 (CXCL10) ELISA kit (Abcam #ab214563), mouse MCP5 (CCL12) ELISA kit (Abcam #ab100723) and mouse/rat CCL2/JE/MCP-1 quantikine ELISA kit (R&D Systems #MJE00). The assays were performed following the manufacturer’s instructions. For BV2 cells, after media aspiration and gentle washing with 1× PBS, a 2.5× loading buffer (Tris HCl 62 mM; 2% sodium dodecyl sulfate; 10% glycerol; 5% β-mercaptoethanol; 0.02 % Bromphenol Blue) was added, and cells were collected. Cells were then sonicated, and the attained protein extracts were processed for western blot. For HMC3 cells, after the media was aspirated, the cells were washed with 1× PBS, trypsinized using 0.25% Trypsin-EDTA (Gibco #25200056), spun down for 5 min at 300 × g. The supernatant was discarded, and the cell pellet was re-suspended in the extraction buffer containing 50mM Tris-HCl, 100 mM NaCl, 5 mM EDTA, and 1 mM EGTA containing the above-mentioned protease and phosphatase inhibitors. Proteins were detected using the following antibodies: rabbit anti-phospho-TBK1/NAK (Ser172) (D52C2); Cell Signaling #5483; 1:1,000); mouse anti-β-actin (Sigma-Aldrich #A2228; 1:2,000); rabbit anti- β-actin (Invitrogen #PA1-183; 1:2000). The following secondary antibodies were used: IRDye® 680RD Goat anti-Rabbit IgG (LI-COR #926-68071; 1:5,000); Goat anti-rabbit StarBright Blue 700 (Bio-Rad #12004162; 1:1,000). Membranes were visualized using the Odyssey CLx LI-COR equipped with the Image Studio software (for BV2) and Bio-Rad ChemiDoc™ MP Imagine System (HMC3). Protein band densitometry was done using the ImageJ software. The target to housekeeping protein (β-actin) expression ratio was used to quantify protein expression, and sham controls were set as 1.

## Assessment of gamma oscillations

Mice were sacrificed 2 weeks, 6 week, 6 months or 1 year post-IR. Animals were anesthetized with isoflurane and decapitated. Brains were quickly dissected and placed in ice-cold artificial cerebrospinal fluid (ACSF) modified for dissection containing (in mM): 80 NaCl, 24 NaHCO_3_, 25 glucose, 1.25 NaH_2_PO_4_, 1 ascorbic acid, 3 Na-pyruvate, 2.5 KCl, 4 MgCl_2_, 0.5 CaCl_2_, 75 sucrose (Sigma-Aldrich) and bubbled with carbogen (95% O_2_ and 5% CO_2_). Horizontal hippocampal sections 3,500 µm thick were prepared from the two hemispheres using a Leica VT1200S vibratome (Leica Microsystems). Slices were immediately placed into an interface holding chamber containing standard ACSF: 124 mM NaCl (Sigma-Aldrich #31434), 30 mM NaHCO_3_ (Sigma-Aldrich #S6014), 10 mM glucose (Sigma-Aldrich #G7021), 1.25 mM NaH_2_PO_4_ (Sigma-Aldrich #S9638), 3.5 mM KCl (Sigma-Aldrich #P9333), 1.5 mM MgCl_2_ (Sigma-Aldrich #2670), 1.5 mM CaCl_2_ (Sigma-Aldrich #C8106) continuously supplied with humidified carbogen gas (95% O_2_ and 5% CO_2_). The chamber was held at 34°C during the slicing process and subsequently allowed to cool down to room temperature (∼22°C) for at least 1 h in order to let the slices recover.

Hippocampal gamma oscillation experiments were performed in a submerged-type recording chamber as described previously ^54^. Local field potentials (LFP) were recorded using microelectrodes filled with standard ACSF placed in the CA3 *stratum pyramidale* and elicited by adding kainic acid (100 nM; Tocris Bioscience) to the extracellular bath.

Oscillation power spectra, fast fourier transformations were obtained from 60 s of LFP recording (segment length 8192 points) using the Axograph X software (Kagi, Berkeley, CA, USA). Frequency variance data was obtained from the power spectra described above using Axograph X. Gamma power was calculated by integrating the power spectral density from 20 to 80 Hz using Clampfit 11.2.

## Statistical analysis

Statistical analyses were performed using GraphPad Prism (GraphPad, Inc., San Diego, CA, USA). Data were presented as mean ± SEM. The unpaired student’s t-test was used to compare SH and IR animals at each time point, aiming to eliminate the effect of animal age. Comparisons of multiple variants per time point were performed using a two-way ANOVA with Bonferroni’s *posthoc* for multiple comparisons. Significance was considered when *p* < 0.05. The statistical analyses, number of animals and *in vitro* experiments applied were noted in each figure legend. For the bulk RNA-seq data analyses, Qlucore Omics Explorer 3.2 software (Qlucore, Lund, Sweden) was used. The GSEAP reranked tool was used for the GSEA analysis, while for single-cell RNA-seq data analyses, both R software (versions 4.3.0 and 4.3.3) and the Seurat package (version 4.0.0) were used.

## Resource availability

Further information and requests for resources and reagents should be directed to and will be fulfilled by the lead contact

---

**Supplementary figure 1.**
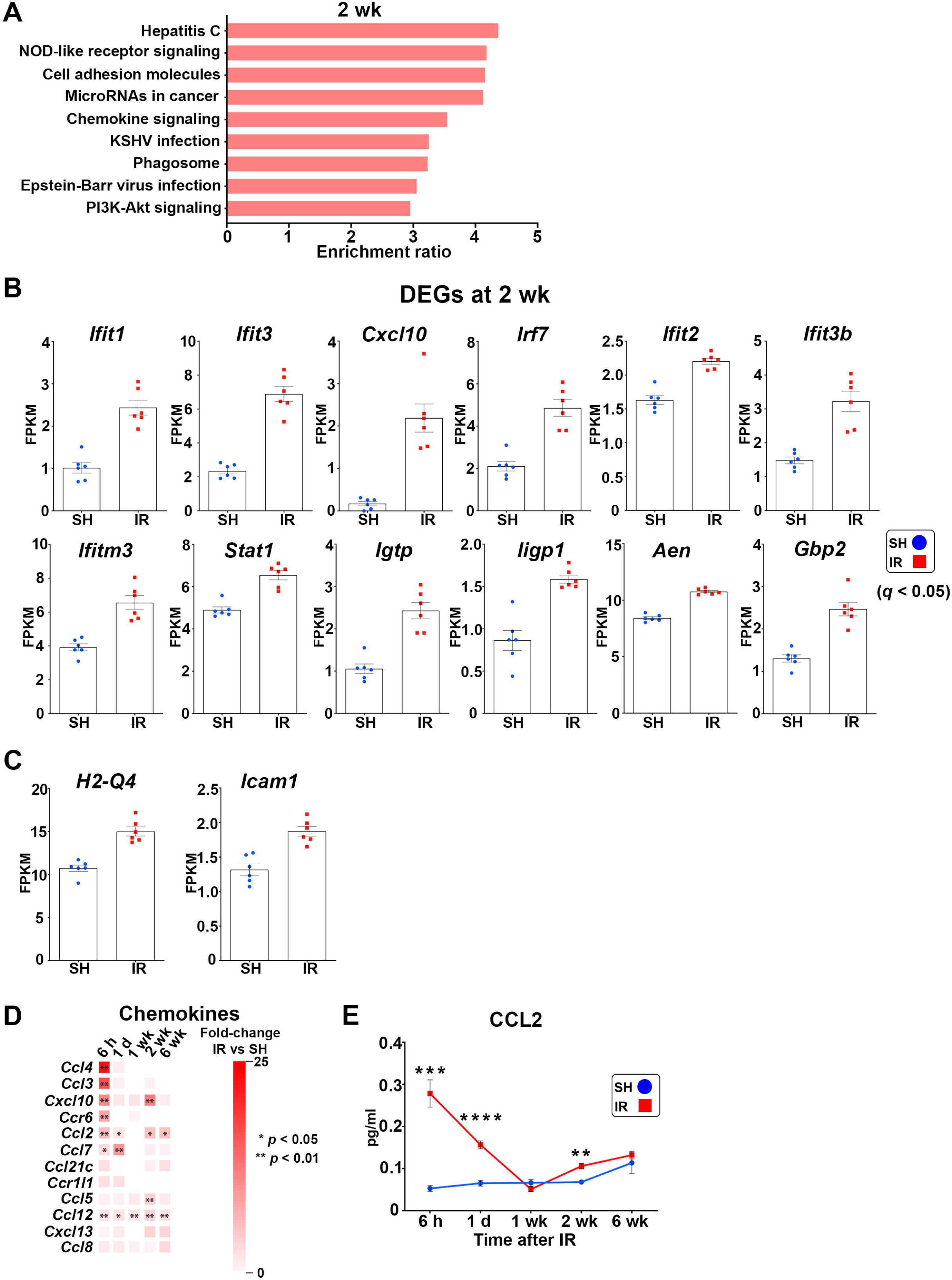
: Irradiation causes biphasic inflammatory waves in the hippocampus. Related to Figure 1. (A) Bar plot showing significantly enriched pathways (*p* < 0.05) 2 wk post-IR revealed by gene set enrichment analysis (GSEA). **(B and C)** Bar plots showing expression of 14 inflammation-related genes in the hippocampus 2 weeks (wk) post-IR (*q* < 0.05), 12 belonging to IFN signaling pathways (**B**). FPKM = fragments per kilobase of transcript per million fragments mapped. (D) The heatmaps depict profiling of chemokines and chemokine receptors in the hippocampus across the time points post-IR. SH, n = 6; IR, n = 6, per time point. *\*p* < 0.05, *\*\*p* < 0.01. (E) Line graphs showing ELISA measurements of CCL2 in the hippocampus across the time post-IR. SH, n = 3-5; IR, n = 3-5. Mean ± SEM, unpaired *t*-test per time point. *\*\*p* < 0.01, *\*\*\*p* < 0.001, *\*\*\*\*p* < 0.0001.

**Supplementary figure 2.**
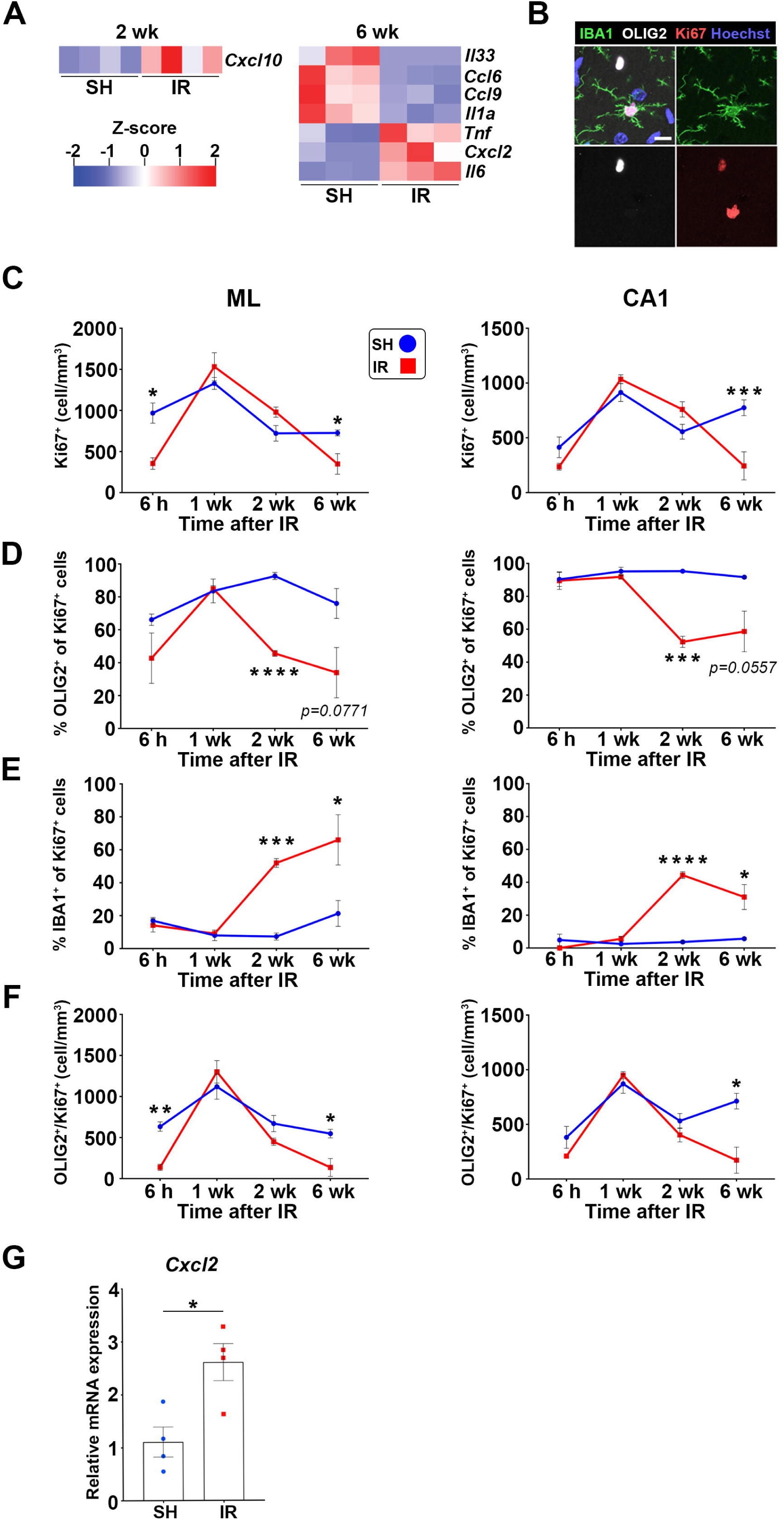
: Delayed microglial responses occurring at later phases post- irradiation. Related to Figure 2. (A) Heatmaps depict significantly regulated genes related to cytokines and chemokines in isolated microglia 2 wk (left) and 6 wk (right) post-IR. (B) Representative confocal images displaying co-labeling of microglia (IBA1+ in green) and oligodendrocytes progenitors (OLIG2^+^; white) with the cell cycle marker Ki67 (red) in the molecular layer (ML) of the hippocampus 2 wk post-IR. Hoechst (blue), nuclear counter stain. Scale bar = 10 μm. (C) Line graphs showing the quantification total of Ki67^+^ cells in the ML (left) and *cornu ammonis* 1 (CA1) region (right) across the time points post-IR. SH, n = 3-4; IR, n = 3-4. Mean ± SEM, unpaired *t*-test per time point. *\*p* < 0.05, *\*\*\*p* < 0.001. (D) Line graphs showing the percentage of OLIG2^+^ cells of total Ki67^+^ cells in the ML and CA1, across the time points post-IR. SH, n = 3-4; IR, n = 3-4. Mean ± SEM, unpaired *t*-test per time point. *\*\*\*p* < 0.001, *\*\*\*\*p* < 0.0001. (E) Line graphs showing percentage IBA1^+^ cells of total Ki67^+^ cells in the ML and CA1 across the time points post-IR. SH, n = 3-4; IR, n = 3-4. Mean ± SEM, unpaired *t*-test per time point. *\*p* < 0.05, *\*\*\*p* < 0.001, *\*\*\*\*p* < 0.0001. (F) Line graphs showing quantification of the proliferating oligodendrocyte progenitors (OLIG2^+^/Ki67^+^) in the ML and CA1 across the studied time points post-IR. SH, n = 3-4; IR, n = 3-4. Mean ± SEM, unpaired *t*-test per time point. *\*p* < 0.05, *\*\*p* < 0.01. (G) Bar plot showing qPCR analysis of proinflammatory chemokine *Cxcl2* in the hippocampus 6 wk post-IR. SH, n = 4; IR, n = 4. Mean ± SEM, unpaired *t*-test. *\*p* < 0.05.

**Supplementary figure 3.**
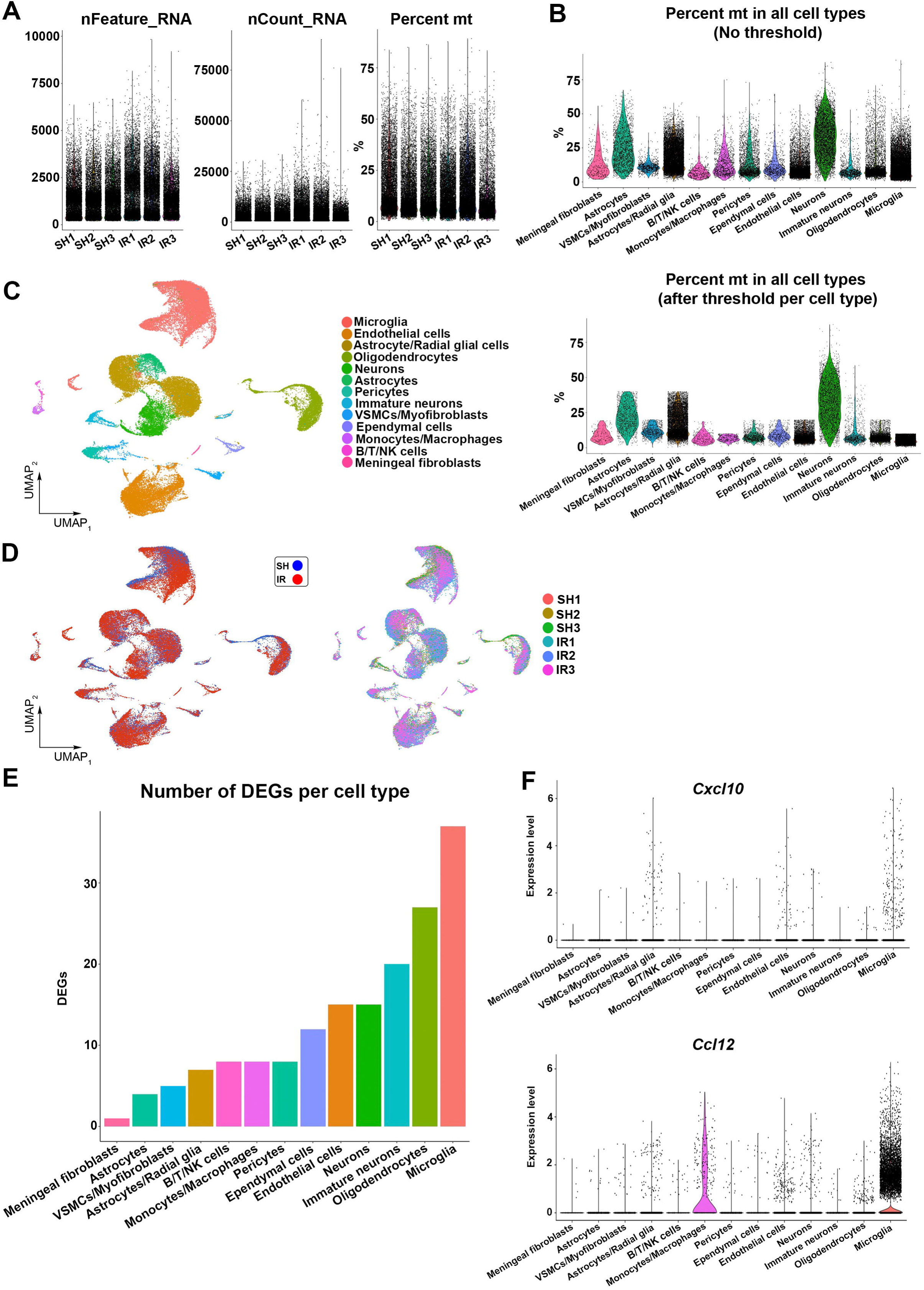
: Processing of the scRNA-seq 2 wk post-irradiation. Related to Figure 3 (A) Violin plots showing the number of detected genes per cell (Left: nFeature), number of reads per cell reflecting the sequencing depths (Middle: nCount), and percentage of the mitochondrial counts per cells (Right: percent mt) after quality control and filtering of doublets in the scRNA-seq datasets obtained from three technical replicates of SH and IR animals. (B) Violin plots showing the percentage of mitochondrial counts per captured cell type before (top) and after (bottom) setting a threshold for inclusion in the analysis. (C) Uniform manifold approximation and projection (UMAP) showing clustering of the cell types captured using our cell isolation protocol from SH and IR hippocampi 2 wk post-IR. N = 3 per group. VSMCs = Vascular smooth muscle cells. B/T/NK cells = B cells, T cells, and natural killer cells, respectively. (D) Left: UMAP showing the captured cell types in SH (blue) and IR (red) animals. Right: UMAP showing that the captured cell types were represented in all three sequenced replicates of the SH and IR animals. (E) Bar plot showing the number of DEGs in all captured cell types. (F) Violin plots showing expression of *Cxcl10* (top) and *Ccl12* (bottom) per cell type post-IR.

**Supplementary figure 4:**
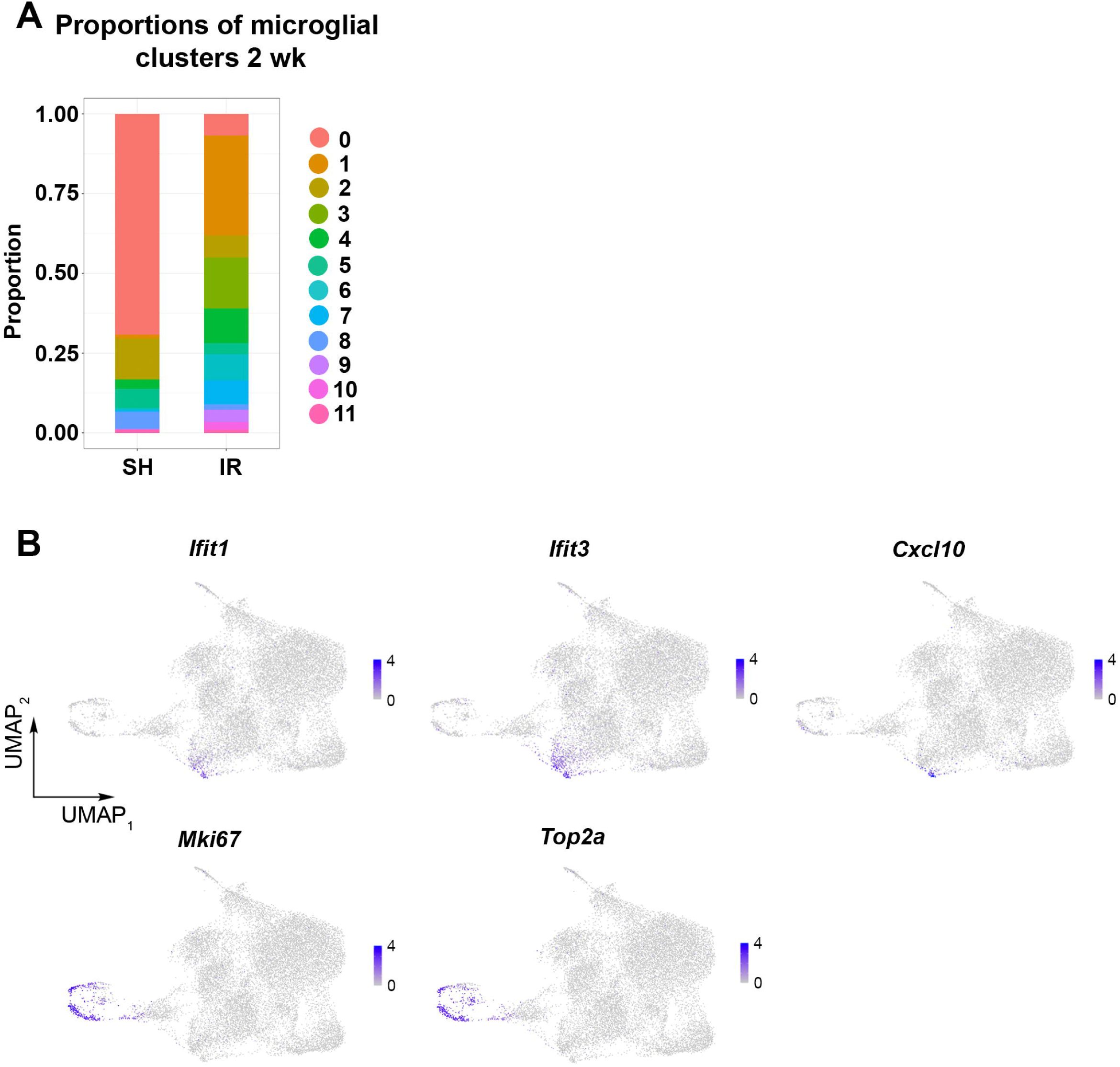
Proportions of microglial clusters 2 wk post-irradiation and expression of genes related to IFN signaling and cell cycle in radiation-associated microglia (RAMs). Related to Figure 3 (A) Stack plot showing the proportions of microglial clusters 2 wk post-IR. (B) UMAPs showing that expression of the IFN-related genes *Ifit1*, *Ifit3,* and *Cxcl10* mainly was in cluster 6, and the expression of the cell cycle genes *Mki67* and *Top2a* was in clusters 7, 9, and 10 of RAM 2 wk post-IR.

**Supplementary figure 5:**
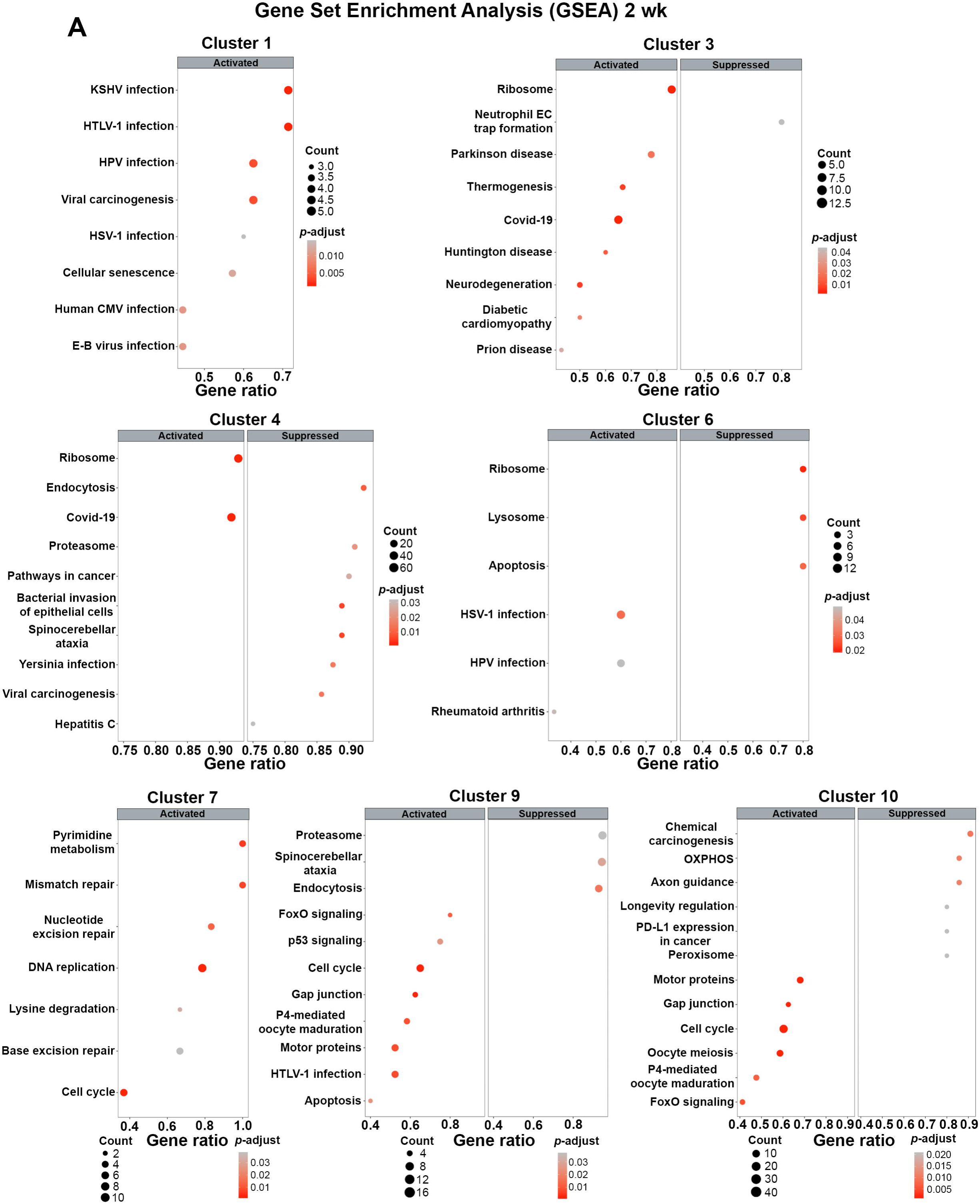
Enriched pathways in RAM 2 wk post-irradiation. Related to Figure 3. (**A**) Dot plots showing the activated and suppressed pathways in the RAM clusters 2 wk post- IR revealed by GSEA.

**Supplementary figure 6.**
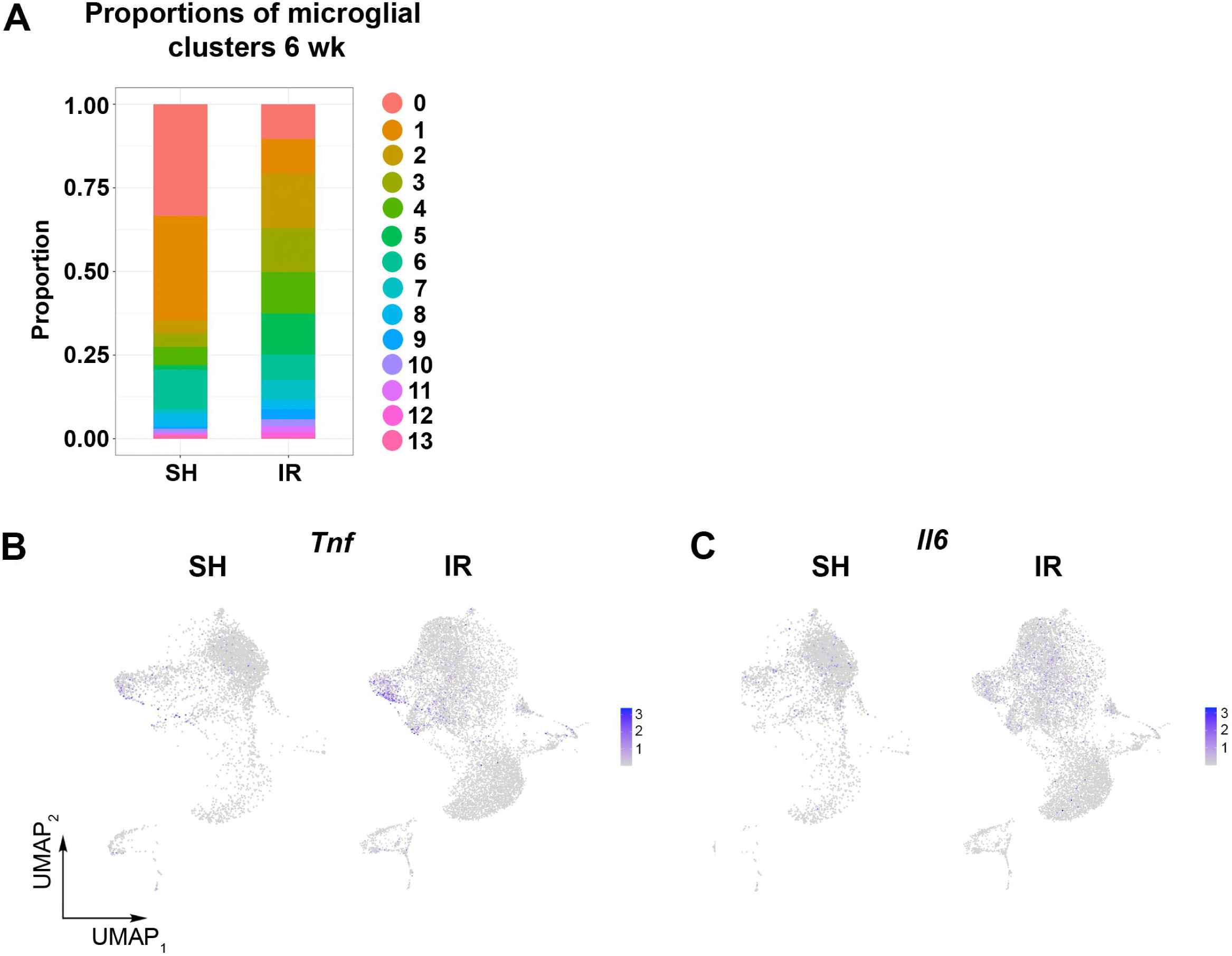
: Proportions of microglial clusters 6 wk post-irradiation and expression of genes *Il6* and *Tnf* in RAMs. Related to Figure 3 (**A)** Stack plot showing the proportion of microglial clusters 6 wk post-IR. (**B** and **C)** UMAPs showing expression of *Tnf* and *Il6* in RAM 6 wk post-IR.

**Supplementary figure 7.**
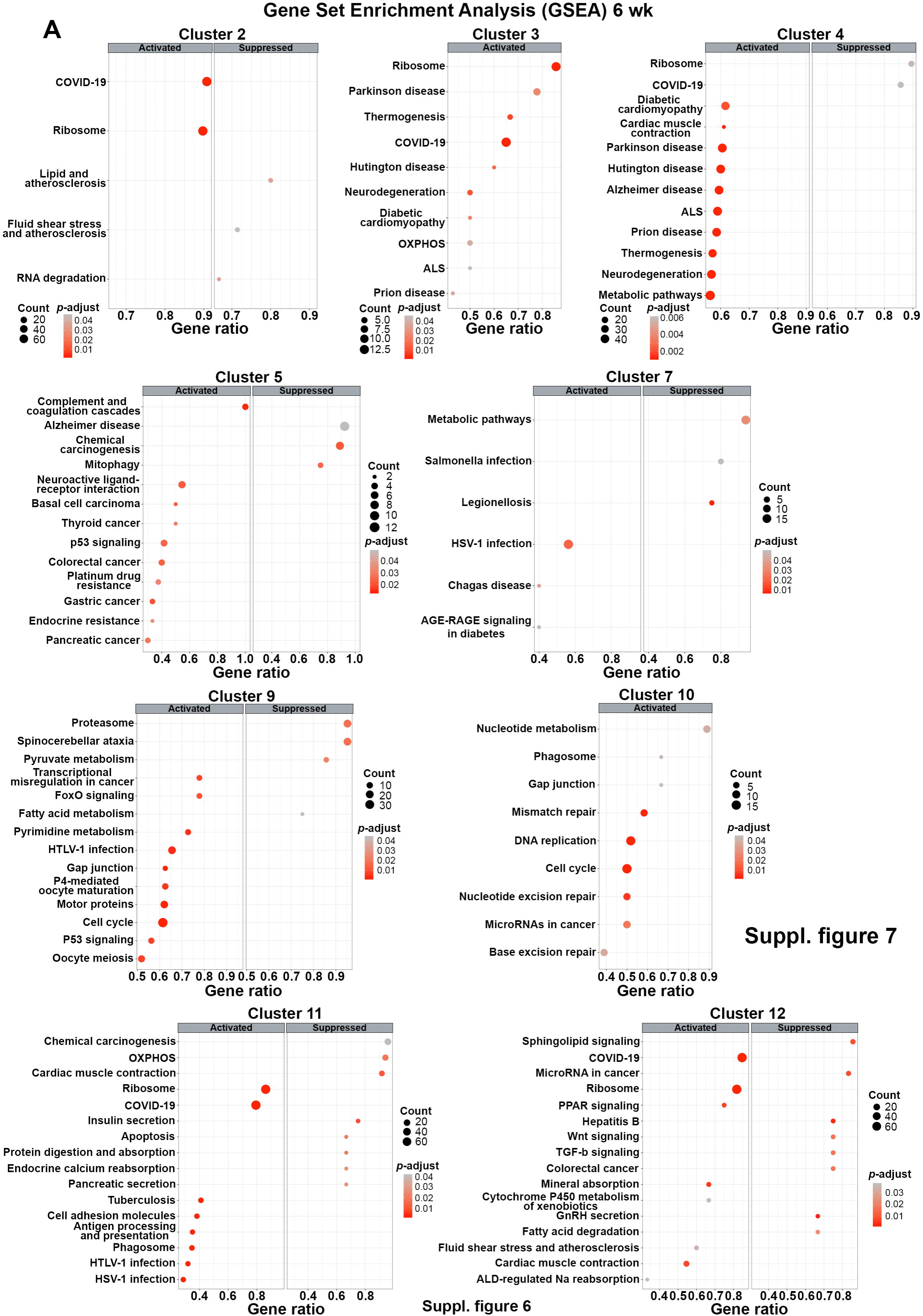
: Enriched pathways in RAM 6 wk post-irradiation. Related to Figure 3. (**A**) Dot plots showing the activated and suppressed pathways in the RAM clusters 6 wk post- IR revealed by GSEA.

**Supplementary figure 8.**
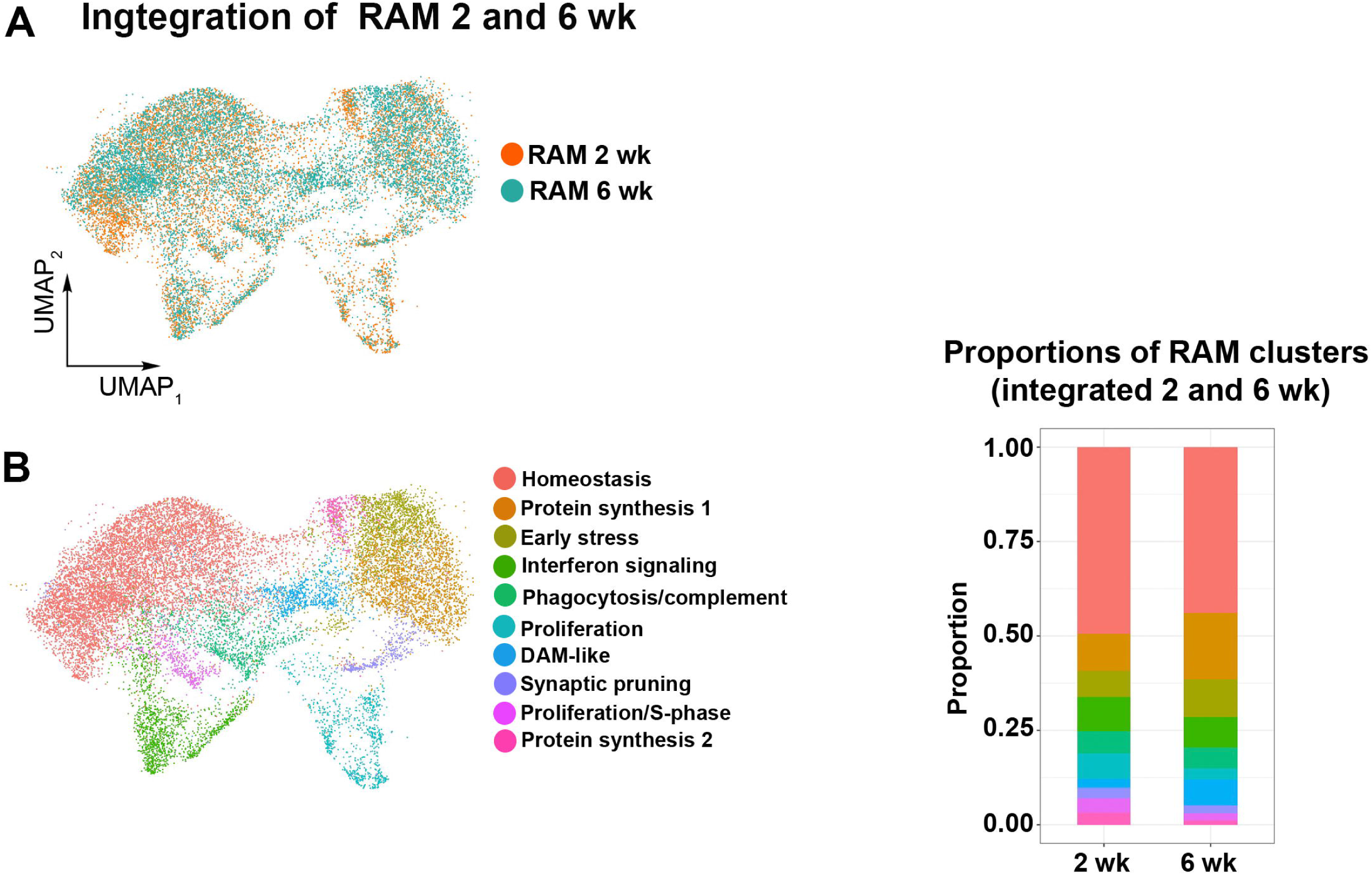
: Integration of the 2 and 6 wk post-irradiation scRNA-seq datasets. Related to Figure 3. (A) UMAP showing integration of 2 and 6 wk post-IR scRNA-seq datasets. (B) Left: Sub-clustering of the integrated cells annotated based on the DEGs. Right: Stack plot showing the proportion of each cluster.

**Supplementary figure 9.**
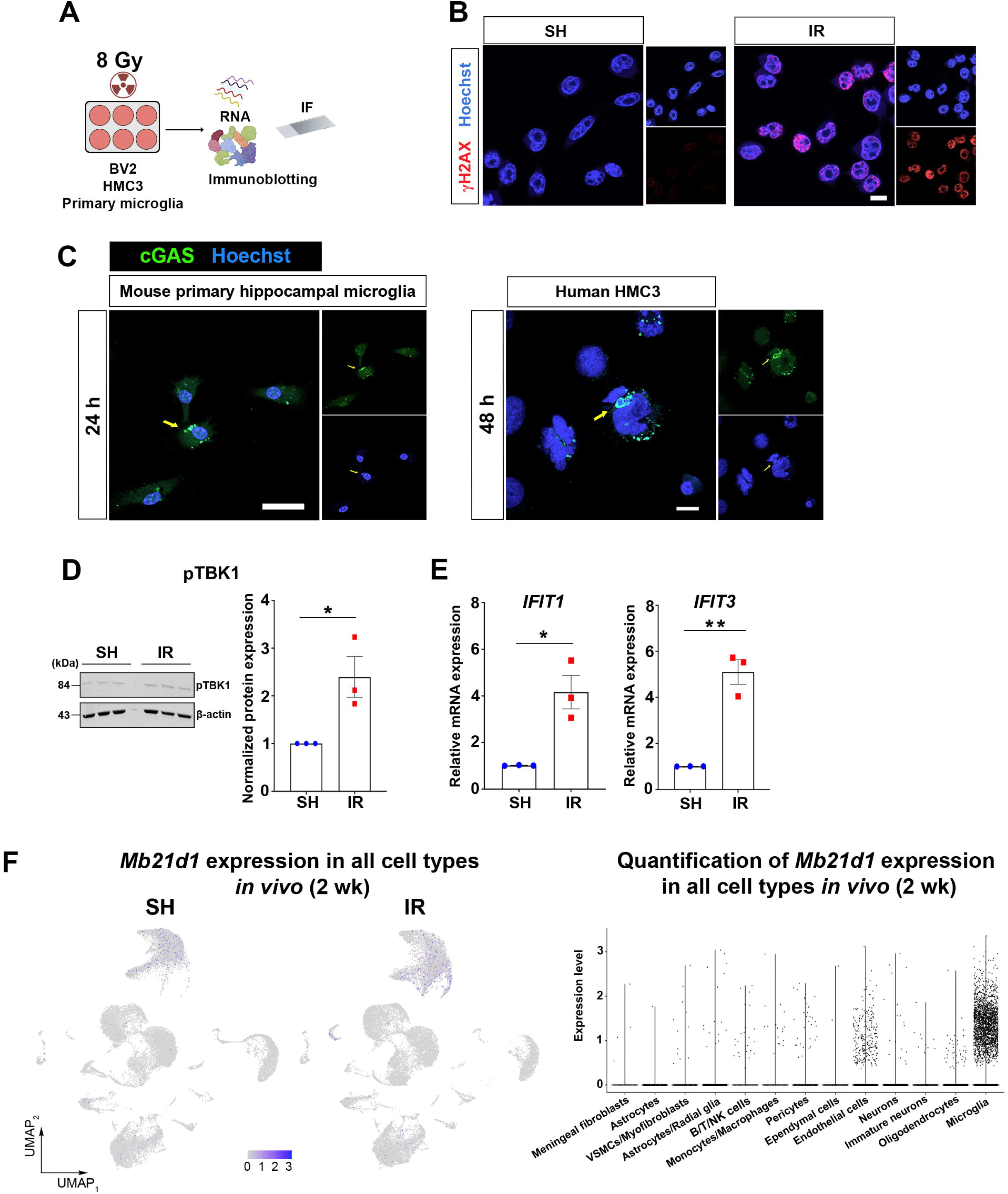
: The role of the cytosolic DNA sensing on the IFN response post-irradiation. Related to Figure 4. **(A)** *In vitro* experimental design. IF = immunofluorescence. **(B)** Representative confocal image displaying expression of the DNA damage response marker γH2AX (red) in the nuclei (Hoechst, blue) of BV2 microglial cells 1 h post-IR. Scalebar = 10 μm. **(C)** Representative immunofluorescence images displaying the micronuclei and cGAS expression (green; indicated by yellow arrows) in mouse primary hippocampal microglia 24 h post-IR (left) and human microglial cell line HMC3 48 h post-IR (right). Hoechst (blue), nuclear counter stain. Scale bar = 20 μm. **(D)** Left: Immunoblots showing the expression of phosphorylated TBK1 (pTBK1) and β- actin expression in HMC3 cells in three technical replicates of SH and IR 48 h post-IR from one experimental set. Right: quantification of pTBK1 48 h post-IR from three independent experiments. Mean ± SEM, unpaired *t*-test. *\*p* < 0.05. **(E)** Bar plots showing qPCR analyses of IFN-related genes *IFIT1* and *IFIT3* in the HMC3 cells 48 h post-IR. Three independent experiments. Mean ± SEM, unpaired *t*-test. *\*p* < 0.05, *\*\*p* < 0.01. **(F)** Left: UMAPs showing expression of *Mb21d1* (encoding cGAS) in all captured cell types, both SH and IR animals, 2 wk post-IR. Right: Violin plot showing quantification of *Mb21d1* expression across cell types.

**Supplementary figure 10:**
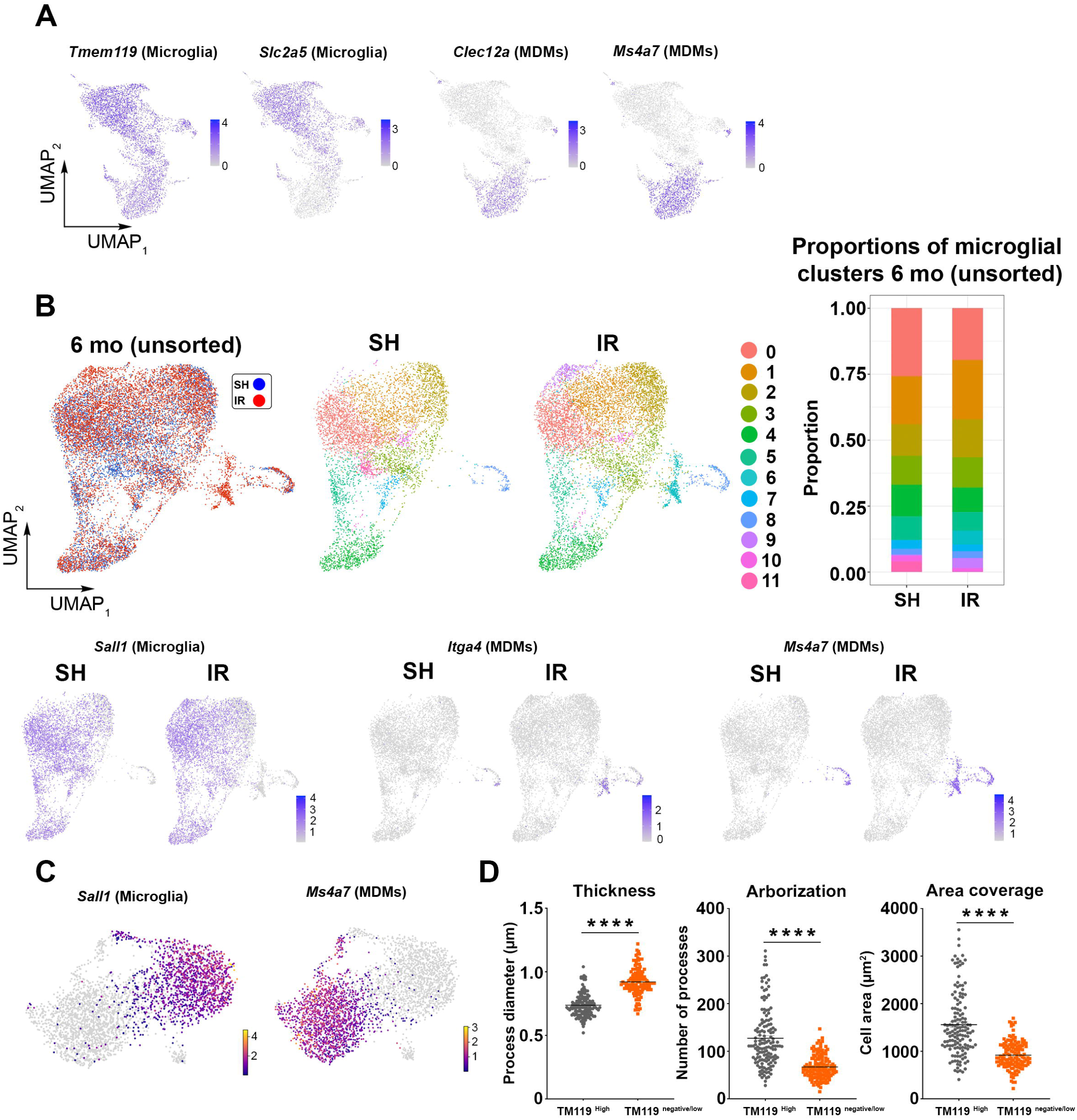
Microglial loss and MDM infiltration into the hippocampus. Related to Figure 5. **(A)** UMAPs showing the expression signature genes of microglia (*Tmem119* and *Slc2a5*) and MDMs (*Clec12a* and *Ms4a7*) in RAM 6 mo post-IR. **(B)** Upper panel: The UMAPs show clustering of unsorted hippocampal microglia from SH (blue) and IR (red) 6 mo post-IR. Sub-clustering revealed 12 clusters. The stack plot shows the proportions of microglial clusters. Lower panel: UMAPs showing the expression of *Sall1* (microglia), *Itga4* (MDMs), and *Ms4a7* (MDMs). Cells in cluster 6 represent MDMs. **(C)** UMAPs displaying expression of the microglial signature gene *Sall1* and MDM signature gene *Ms4a7 in* RAM 6 mo post-IR where RNA velocity analysis was applied. **(D)** Dot plots showing morphological comparisons between the TMEM119^high^ and TMEM119^low^ cells 6 mo post-IR. n = 144 - 229 cells collected from n = 3 animals per group. Mean ± SEM, unpaired *t*-test. *\*\*\*\*p* < 0.0001. TM119 = TMEM119.

**Supplementary Figure 11.**
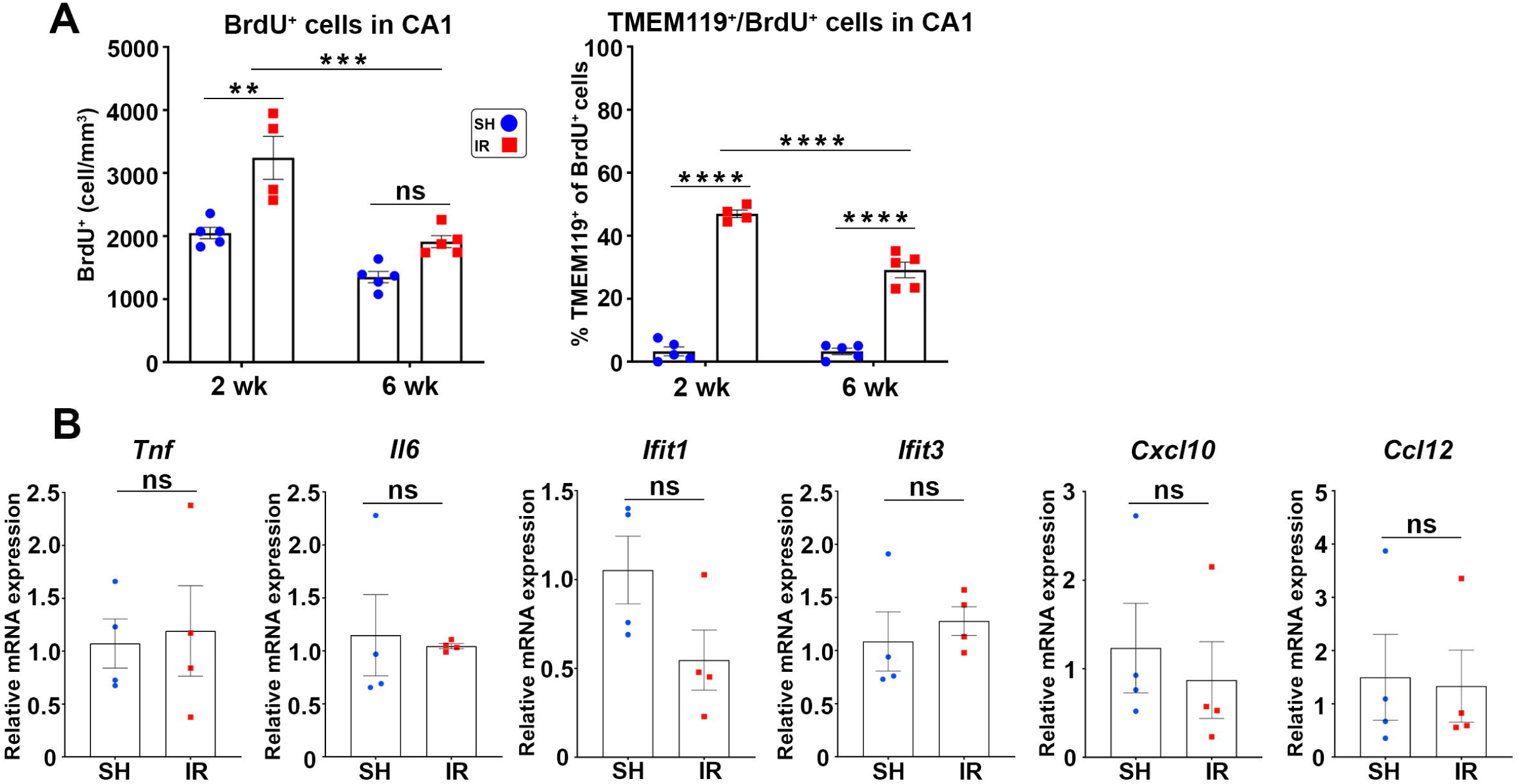
: Quantification of BrdU and the expression of inflammatory mediators 6 mo post-irradiation. Related to Figure 5. (A) Bar plots showing quantification of total BrdU^+^ cells (left) and percentage of TMEM119^+^ of total BrdU^+^ cells (right) in the CA1 2 and 6 wk post-IR. 2 wk, n = 4-5; 6 wk, n = 5. Mean ± SEM, two-way ANOVA with Bonferroni’s *post hoc* test for multiple comparisons. *\*\*p* < 0.01, *\*\*\*p* < 0.001, *\*\*\*\*p* < 0.0001. ns = not significant. (B) Bar plots showing qPCR analyses of proinflammatory mediators in the hippocampus 6 mo post-IR. SH, n = 4; IR, n = 4. Mean ± SEM, unpaired *t*-test. ns = not significant.

**Supplementary figure 12.**
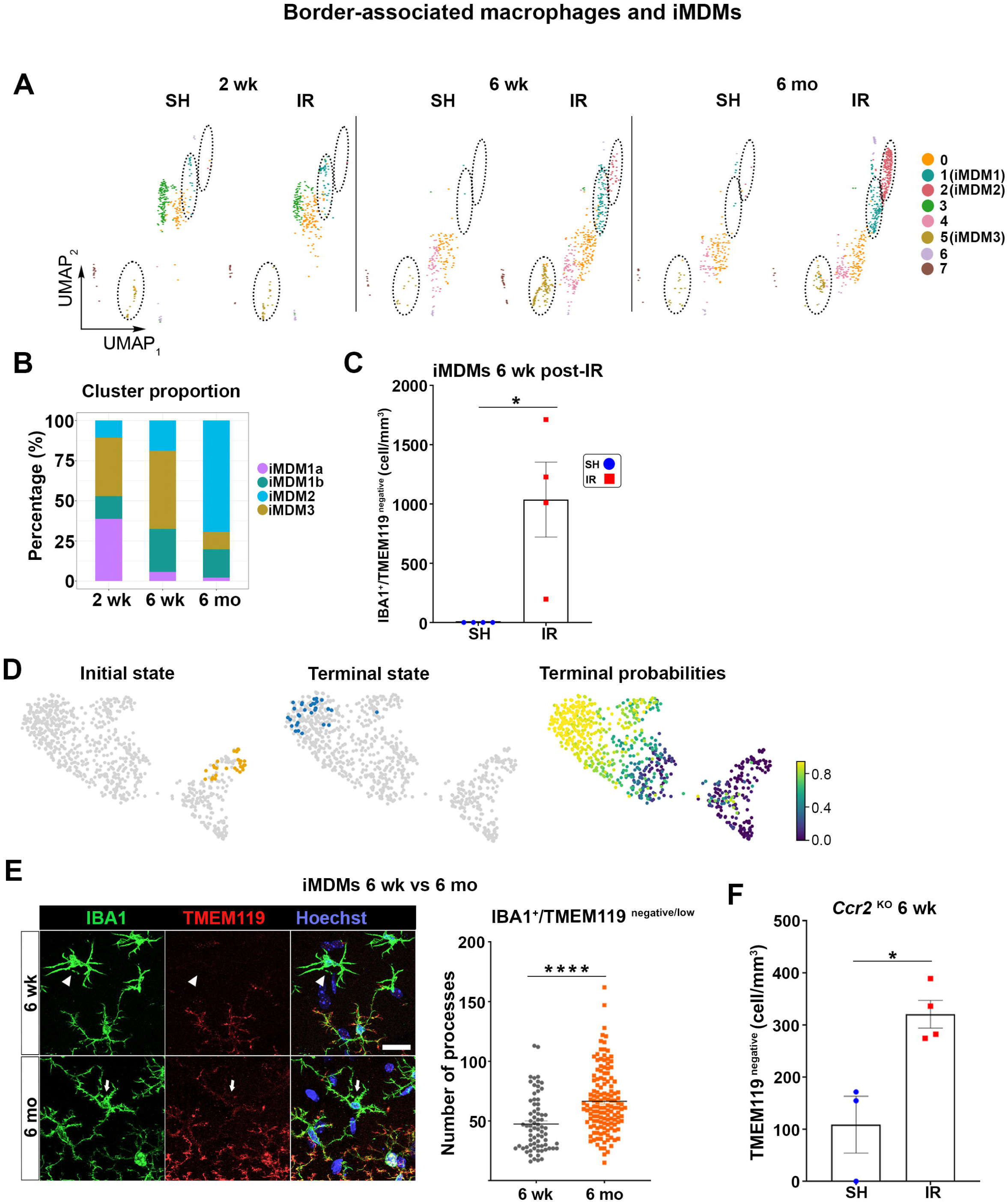
: MDM dynamics over time post-irradiation. Related to Figure 6. (A) UMAP showing the macrophage populations detected in the hippocampus of SH and IR animals. IR-induced MDMs (iMDMs) were cluster 1 (= iMDM1 in Figure 6A), 2 (= iMDM2), and 5 (= iMDM3). (B) Stack plot showing the proportion of iMDM clusters over time post-IR. (C) Bar plot showing quantification of IBA1^+^/ TMEM119^negative^ in the CA1 6 wk post-IR. SH, n = 4; IR, n = 4. Mean ± SEM, unpaired *t*-test. *\*p* < 0.05. (D) UMAPs visualizing the probabilities of the initial and terminal states among the iMDMs. (E) Comparisons between the IBA1^+^/TMEM119^negative^ ^and^ ^low^ cells detected 6 wk and 6 mo post-IR. Left: Representative confocal images displaying co-labeling of IBA1^+^ cells (green) TMEM119^+^ (red) in the CA1 at both time points. Hoechst (blue), nuclear counter stain. Arrows indicate TMEM119^low^, and arrowheads indicate TMEM119^negative^. Scale bar = 10 μm. Left: Quantification of the number of processes per cell. n = 74 -141 cells collected from n = 3 - 4 animals per group. Mean ± SEM, unpaired *t*-test. *\*\*\*\*p* < 0.0001. (F) Bar plot showing quantification of IBA1^+^/ TMEM119^negative^ in CA1 of *Ccr2* knockout (KO) mice 6 wk post-IR. SH, n = 3; IR, n = 4. Mean ± SEM, unpaired *t*-test. *\*p* < 0.05.

**Supplementary figure 13.**
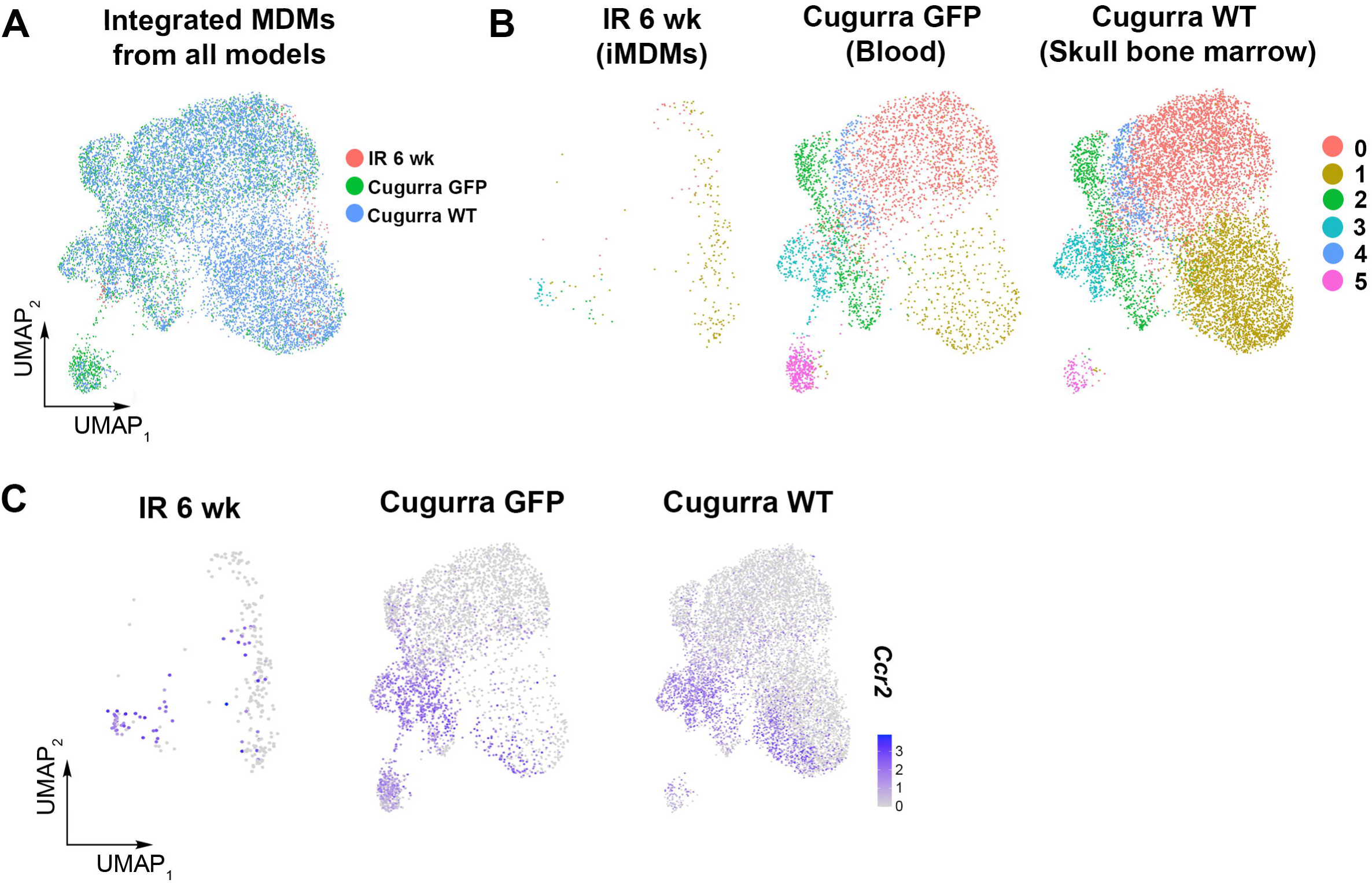
: Comparison of iMDMs with blood or CNS-associated bone marrow-derived MDMs. Related to Figure 6. (A) UMAP showing embedding of the 6-wk iMDMs post-IR with the monocyte, macrophage, and microglia populations presented in scRNA-seq from (Cugurra *et al.*, 2021) derived from the blood (Cugurra GFP) or CNS-associated bone marrow (Cugurra WT). **B**) Sub-clustering of the cells in (**A**). (**C**) Visualization of *Ccr2* expressing cells in each population.

**Supplementary figure 14.**
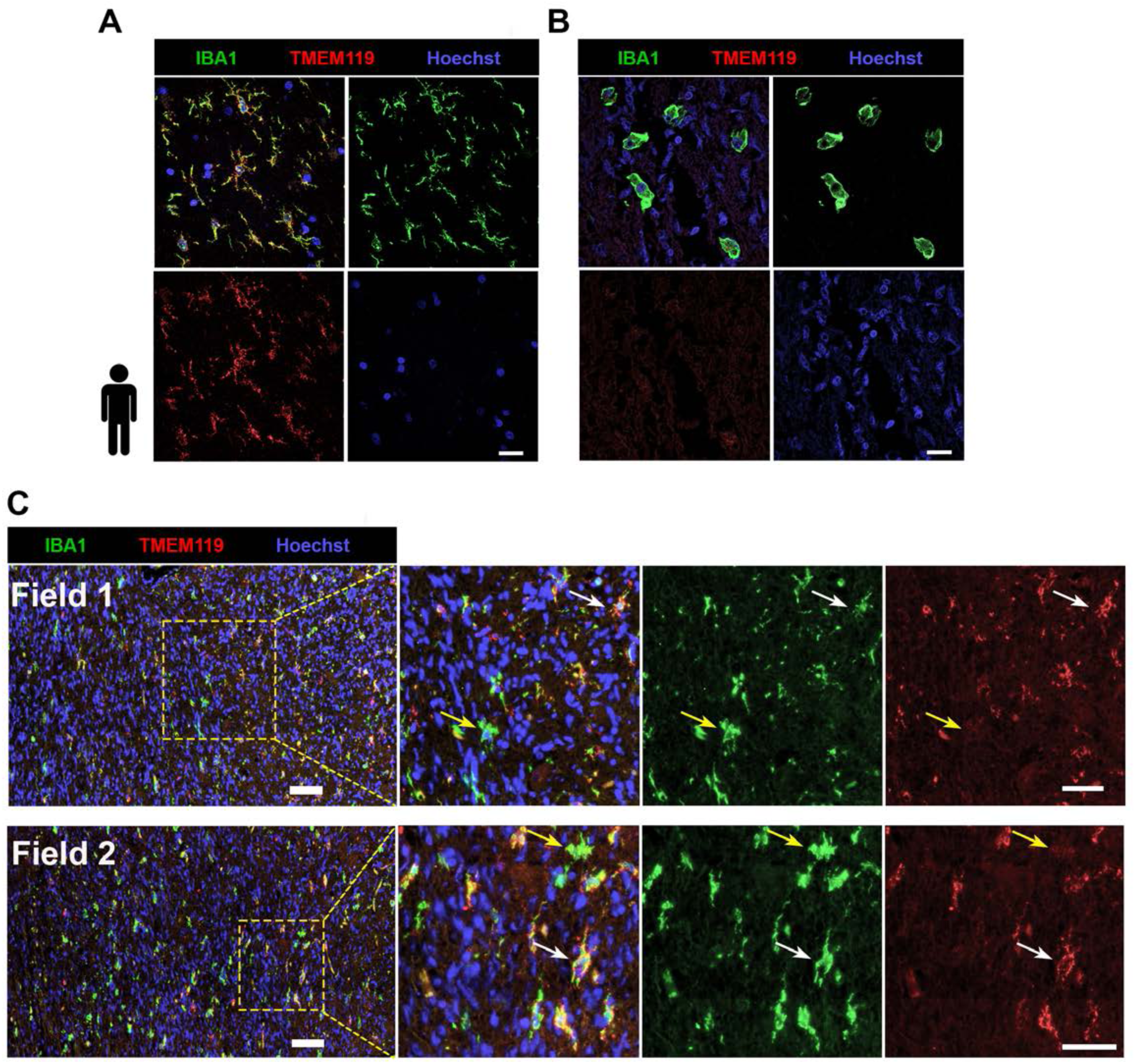
: Immunofluorescence staining of human samples. Related to Figure 6. **(A** and **B**) Representative confocal images showing the specificity of the immunofluorescence stainings for expression of IBA1 (green) and TMEM119 (red) cells in human brain tissues. (**A**) Showing colocalization of IBA1 and TMEM119 in postmortem hippocampal tissue donated by a 54-year-old healthy male (used as positive control for both stainings). (**B**) Showing IBA1^+^ /TMEM119^negative^ macrophages infiltrating a surgically resected ependymoma tissue (used as a positive control for IBA1 and a negative control for TMEM119). Hoechst (blue), nuclear counter stain. Scale bar = 20 μm. (C) Representative images displaying the expression of IBA1 (green) and TMEM119 (red) in cancer-free hippocampus of a medulloblastoma patient 5 years post-radiotherapy depicted from two independent fields. Yellow arrows indicate IBA1^+^/TMEM119^negative^ cells; white arrows indicate IBA1^+^/TMEM119^+^ cells. Hoechst (blue), nuclear counter stain. Scale bar = 100 μm (overview) and = 50 μm (closeup).

**Supplementary figure 15.**
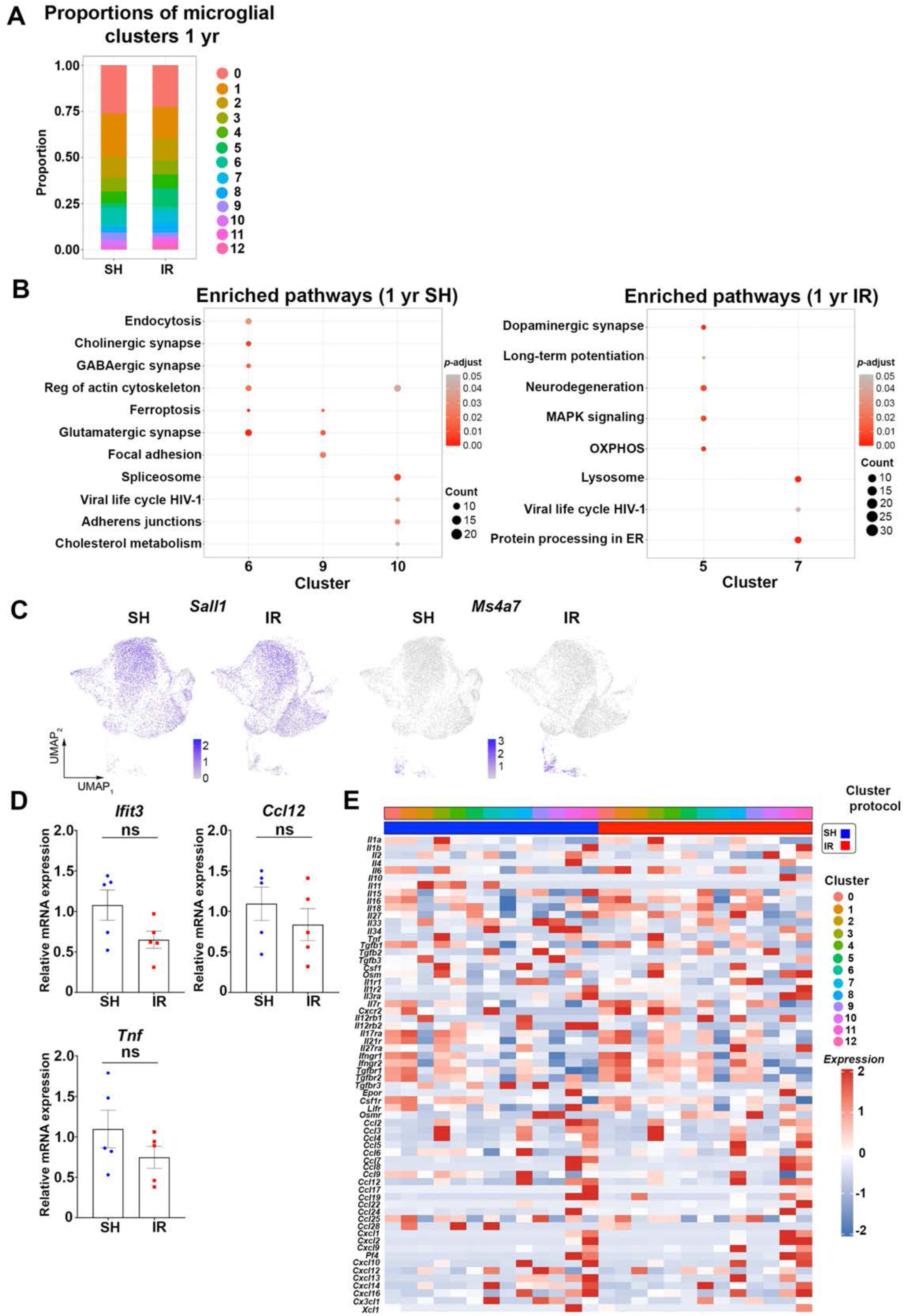
: Microglial alterations and the inflammatory response 1 year (yr) post-irradiation. Related to Figure 6. (A) Stack plot showing the proportions of microglial clusters 1 yr post-IR. (B) Dot plot showing the enriched pathways 1 yr post-IR in SH (Left) or RAM (right) revealed by GSEA. (C) UMAPs showing expression of the microglia signature genes *Sall1* and the MDM signature gene *Ms4a7*. (D) Bar plots showing qPCR analyses of inflammatory mediators *Ifit3*, *Ccl12,* and *Tnf* in the hippocampus 1 yr post-IR. SH, n = 5; IR, n = 5. Mean ± SEM, unpaired *t*-test. ns = not significant. (E) Heatmap showing average expression of the cytokines, chemokines, and their receptors detected in each microglial cluster from SH and RAM 1 yr post-IR.

**Supplementary figure 16.**
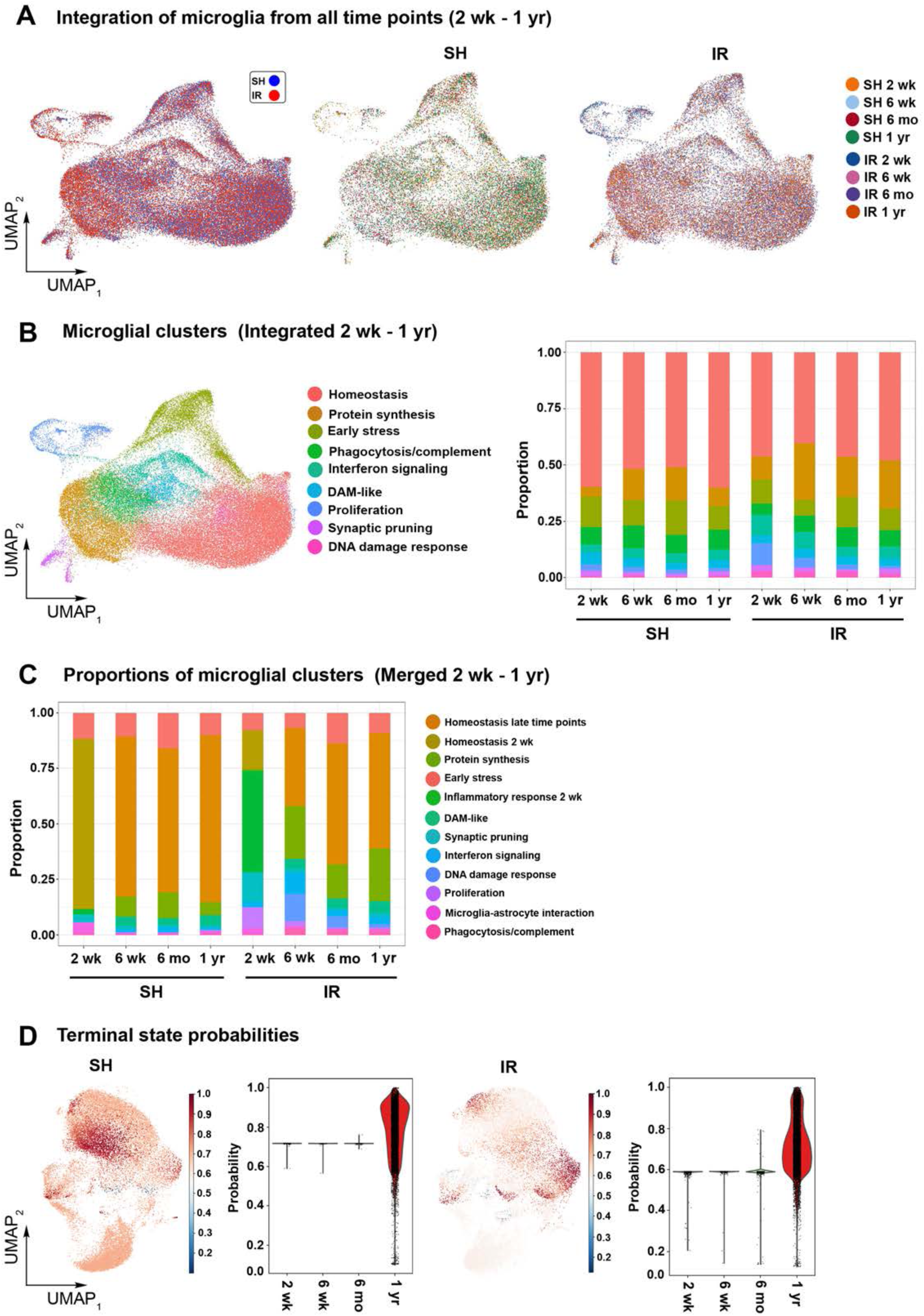
: Longitudinal overview of RAM (2 wk - 1 yr). Related to Figure 7 (A) Left: UMAP showing integration of RAM and their age-matched SH control microglia obtained from the 2-wk to 1-yr time points. Right: SH and RAM were split and annotated by time point. (B) Left: Sub-clustering of the integrated microglia obtained from the 2-wk to 1-yr time points. Right: the proportion of each cluster over the studied time points. Clusters were annotated based on the DEGs. (C) The proportions of microglial clusters after merging the dataset obtained from the 2-wk to 1-yr time points. Clusters were annotated based on the DEGs. (D) UMAPs visualizing the terminal state probabilities in SH and RAM over the studied time points. The violin plots show terminal state probabilities across the studied time points.

## Notes

### Summary of Updates

Extra in vitro work is added, and the single-cell and computational analyses have been amended.

